# Genetic diversity and fine-scale structure of two populations of a subterranean rodent, *Ellobius talpinus*

**DOI:** 10.1101/2024.10.16.618669

**Authors:** Anna I. Rudyk, Kristina V. Kuprina, Arman M. Bergaliev, Svetlana A. Galkina, Anna E. Romanovich, Eugene A. Novikov, Elena V. Volodina, Antonina V. Smorkatcheva

## Abstract

Subterranean rodents represent an interesting model system for molecular ecologists. Their lifestyle is associated with fragmented environments, limited dispersal ability, and low fecundity. These characteristics all are expected to increase inter- and intrapopulation differentiation and reduce intra- population genetic diversity, yet the published empirical data revealed the lack of generality of this pattern. This emphasizes the importance of accumulating more data on individual species to understand what factors shape the genetic dynamics of populations. This study represents the first characterization of the population genetic diversity and fine-scale genetic structure of a highly specialized subterranean vole, *Ellobius talpinus*. We used nine microsatellite loci and a fragment of the mitochondrial D-loop to investigate genetic patterns of two distant populations: the Novosibirsk population at the extreme northeastern edge of the species range, and the sub-peripheral Saratov population. As a result, the two populations exhibit distinct patterns of genetic structure. The Novosibirsk population showed a low nuclear diversity with an observed heterozygosity (*Ho*) of 0.34 and an unbiased expected heterozygosity (*uHe*) of 0.55. Mitochondrial D-loop was nearly monomorphic with over 90% of the observed haplotypes being identical, resulting in a haplotype diversity (*Hd*) of 0.14 and nucleotide diversity (π) of 0.0004. In contrast, the Saratov population displayed moderate nuclear (*Ho* = 0.64; *uHe* = 0.76) and high mitochondrial variation (*Hd* = 0.83; *π* = 0.034), compared to the surface-dwelling voles. Nine haplotypes representing four well-differentiated mitochondrial clades were found in the Saratov population. In a heterogeneous landscape, significant genetic differentiation was revealed at both intermediate (dozens of kilometers) and fine (several kilometers) scales. However, within the continuous suitable habitat, no fine-scale spatial structure was observed apart from that caused by kin clustering. The spatial genetic patterns revealed in *E. talpinus* appear to reflect a combination of effects of strong social structure, local natural and anthropogenic barriers limiting dispersal opportunities, and occasional long-distance dispersal.

## 1. INTRODUCTION

Revealing the relationship between the dynamics of genetic structure of populations and various external factors and species traits is important for understanding evolutionary trends. It also provides a theoretical basis for effective conservation measures. Subterranean rodents represent an interesting model system for molecular ecologists. Their lifestyle is associated with naturally fragmented environments, limited dispersal ability, and low fecundity which characteristics all are expected to increase inter- and intrapopulation differentiation and reduce intra-population genetic diversity (Nevo 1979 1990; Bush et al. 2000; Steinberg and Patton 2000; De Kort et al. 2021). Empirical evidence appears to support the first prediction (e.g., Daly and Patton 1990; Wlasiuk et al. 2003; Mora et al. 2010; Roratto et al. 2015; Visser et al. 2018; but see Reuber et al. 2024). On the other hand, genetic diversity of subterranean rodents varies across species and populations from the predicted low levels (some species of *Ctenomys* – Lacey 2001; Wlasiuk et al. 2003; El Jundi and de Freitas 2004; Goncalves, de Freitas 2009; *Spalacopus cyanus* - Opazo et al. 2008; Begall and Honeycutt 2022; *Heterocephalus glaber* - Ingram et al. 2015) to high or very high (*Ctenomys flamarioni* - Fernández-Stolz et al. 2007; *Nannospalax leucodon* - Karanth et al. 2004; Popa et al. 2014; *Bathyergus suillus* - Visser et al. 2014; *Fukomys damarensis* - Mynhardt et al. 2021). This diversity emphasizes the importance of accumulating more data on individual species to understand which factors shape the genetic dynamics of populations. The differences in genetic diversity may arise from distinctive habitat configurations and species-specific features such as life histories, breeding systems, or dispersal strategies (Lacey 2000; Begall et al. 2007; Visser et al. 2018; De Kort et al. 2021). At the intraspecies scale, variation of genetic polymorphism may be related to unequal historical or current fragmentation levels, climate or abundance (Karanth et al. 2004; Lin et al. 2008; Ingram et al. 2015; Kittlein et al. 2021). An important factor influencing genetic diversity is the position of the population relative to the center of the species range. Peripheral populations are expected to be small, isolated, and occur in ecologically marginal habitats where selection pressures are likely to be more intense (Lawton 1993). Such populations should have low genetic diversity due to high inbreeding, genetic drift and directional selection (Consuegra et al. 2005; Hampe and Petit 2005; Arnaud-Haond et al. 2006; Eckert et al. 2008; Carvalho et al. 2019). However, reviews of published empirical data (Pironon et al. 2016; De Kort et al. 2021) revealed the lack of generality of this pattern. Thus, the association between central/peripheral position of populations and their genetic polymorphism deserves further examination. Comparison of the genetic structure and diversity of subterranean species or populations that differ with respect to any of the mentioned characteristics can contribute to our understanding of factors affecting population dynamics, viability and extinction risks and the microevolutionary processes. The studies providing estimates based on different types of markers are most informative.

Mole voles (*Ellobius*) are subterranean members of the young and species-rich subfamily Arvicolinae (Fig. 1). This highly specialized myomorph genus provides a unique opportunity to be compared with both ecologically similar hystricognath rodents and closely related non-subterranean voles. The genus *Ellobius* itself is remarkable by dramatic interspecific and intraspecific variation in rates of chromosomal evolution and by the aberrant sex determination systems in most of the species (Lyapunova et al. 1980; Bakloushinskaya 2009; Bakloushinskaya et al. 2010 2019; Romanenko et al. 2019; Tambovtseva et al. 2022). Mole voles are of particular interest to behavioral ecologists because they live in extended cooperative groups, similar to those of some social mole-rats (Shubin 1978; Davydov 1988; Evdokimov 2001). Moreover, the northern mole vole, *E. talpinus,* occupies various habitats and displays inter-population differences in demography and social structure in different parts of its vast range (Evdokimov 2001; Novikov et al. 2007), which provides the opportunity to use intra-species comparison for testing various hypotheses. Taken together, these features make *Ellobius* a unique model system to study population processes, adaptive evolution and speciation. In addition to the theoretical significance, investigation of the mole voles’ population genetics is important for conservation of these rodents which are considered endangered in Ukraine (Akimov 2009). Meanwhile, mole voles apparently remain the least studied truly subterranean rodents in terms of population genetics. Although the variability of *cytochrome B* and a few protein-coding nuclear genes have been evaluated in the context of phylogeography for several *Ellobius* species (Bogdanov et al. 2015; Alireza et al. 2018, Lebedev et al. 2020; Tambovtseva et al. 2022), intrapopulation genetic diversity and genetic structure at local or fine spatial scale have never been explored in these rodents.

**Fig. 1.**
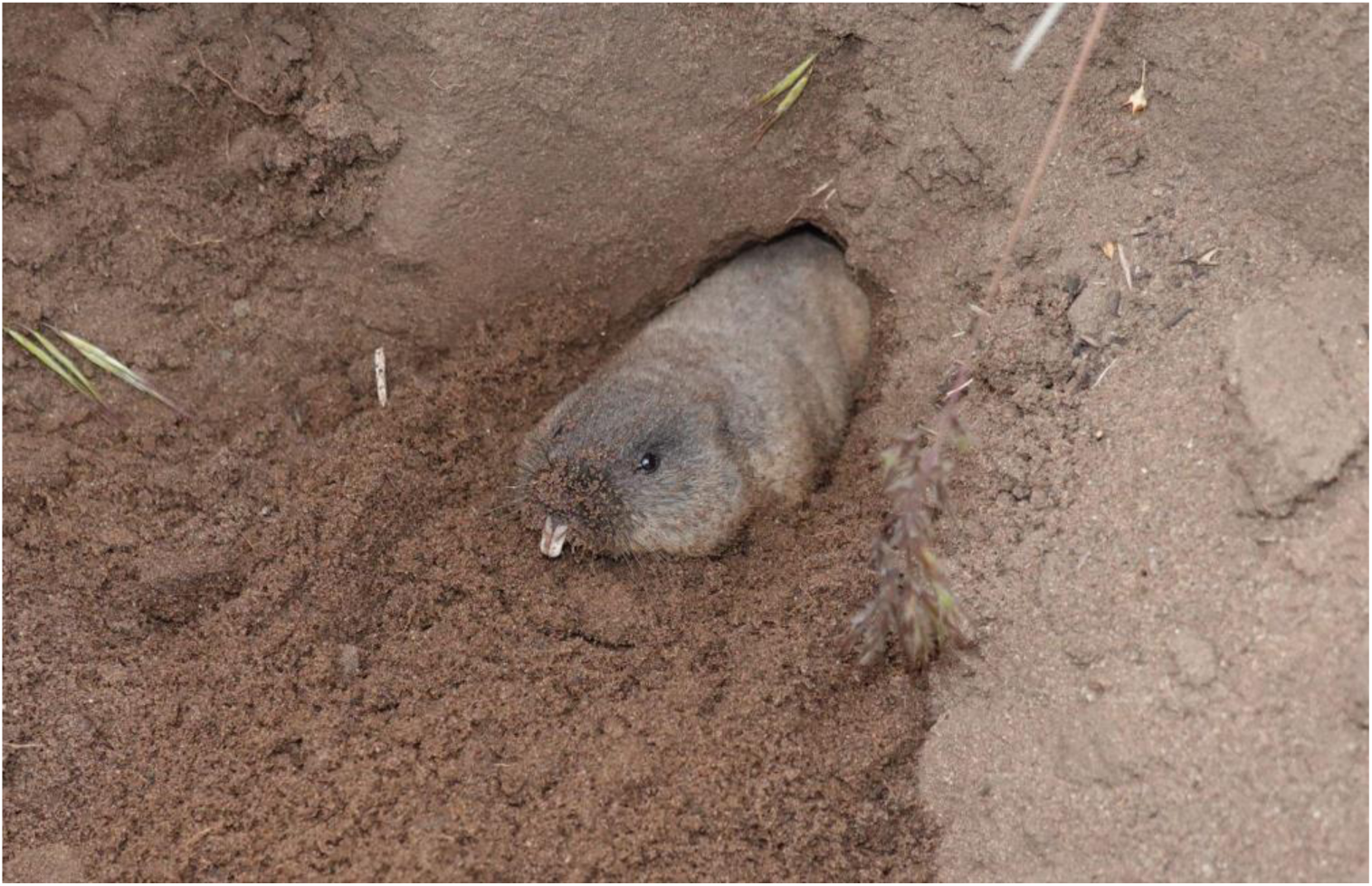
The northern mole vole pushes soil out of its burrow.

In this study, we use a combination of highly variable nuclear (nine microsatellite loci) and mitochondrial DNA (D-loop) markers to quantify, for the first time, the genetic diversity and fine-scale genetic differentiation in *E. talpinus*. We explore and compare the genetic patterns of two populations from distant parts of the species range (Fig. 2A). The first study population is located in the Novosibirsk region, at the extreme northeast of the species’ range, and is isolated from the nearest known localities by the Ob River (Fig. 2B). This area lies in the forest-steppe zone where grassy habitats suitable for mole voles are separated by forests, wetlands and bodies of water causing strong population fragmentation (Kravtsov et al. 2002; Novikov et al. 2007). The second study population is located in the Saratov Region, in the transitional zone between the dry steppes and demi-deserts (Fig. 2C). These are landscapes in which the habitats preferred by *E. talpinus* predominate and where the species reaches its greatest abundance (Popov 1960; Sludsky 1978; Evdokimov 2001). This study aimed to answer the following questions: (i) Do both mole vole populations exhibit reduced genetic polymorphism and strong fine-scale structure, compared to closely related non-subterranean species? (ii) Is genetic diversity lower in the marginal Novosibirsk population than in sub peripheric Saratov population?

**Fig. 2.**
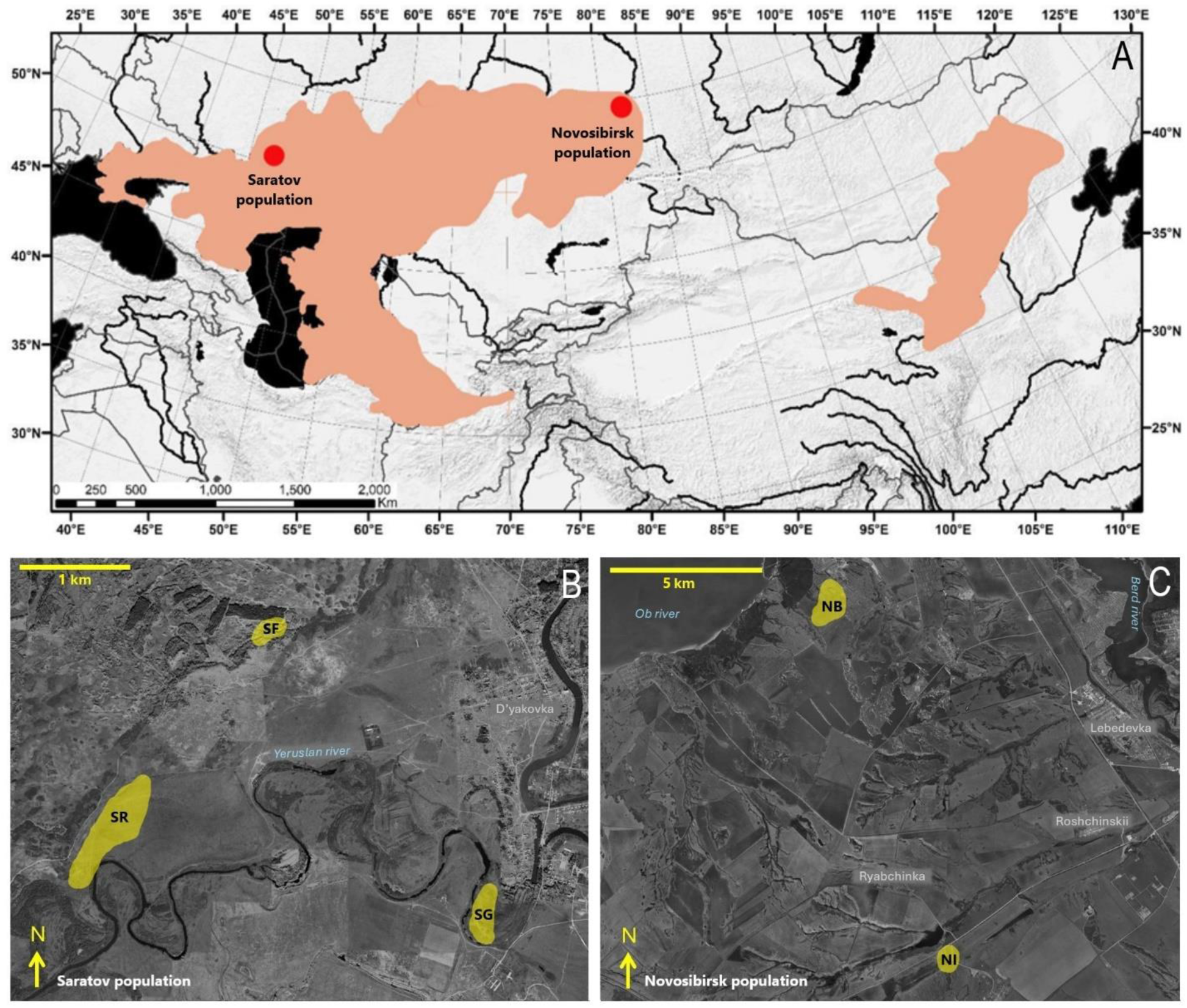
Distribution map of the *Ellobius talpinus* species and geographical locations of two studied populations (red dots) (species’ range is modified from Krystufek and Shenbrot 2022) (A). Satellite maps showing the sampling sites within the Saratov region (B) and Novosibirsk region (C).

## 2. MATERIALS AND METHODS

### 2.1 Study populations and sampling

In the Novosibirsk region, we sampled two sites 13.4 km apart (hereinafter NB and NI; Fig. 2B), both dominated by meadow-steppe associations preferred by mole voles. The area between them was a mosaic of various habitats and included natural and anthropogenic barriers such as rivers, wetlands and intensively used plowed farmland. The trapping areas in NB and NI were approximately 20 and 3 ha, respectively.

In the Saratov Region, the sampling was carried out in three sites (Fig. 2C). Two of them, SR and SF, were within a large continuous area of psammophyte steppe. They were separated by 1.8 km of grassland used for livestock grazing, without any physical barriers. The third site (SG) was 3.2 and 3.3 km from the sites SR and SF, respectively and was separated from them by the bends of the Eruslan River and areas unsuitable for mole voles due to severe overgrazing. It was in an abandoned garden where mole voles inhabited the clearings overgrown by *Krascheninnikovia ceratoides* on dense clay soils. The trapping areas in SR, SF and SG were approximately 25, 3 and 6 ha, respectively.

The animals were captured using Golov’s live-traps (Golov 1954). Precise locations of each trapping point were quantified using a hand-held GPS (± 5 m error), and the burrow (= group) ID were noted. Burrows were considered different if they were separated by a distance of at least 20 m and were free of either old or fresh mounds. Animals were sexed and weighed. To distinguish adults from juveniles, the joint width of both upper incisors was measured by electron callipper to the nearest 0.01 mm in accordance with the method developed for a closely related species, *Ellobus tancrei* (Kuprina and Smorkatcheva 2018). The animals from the Saratov population were tagged with microtransponders 1.25 x 7 mm (Star Security Technologies Co., Shanghai, China) for further identification. From each animal, a non-destructive tissue sample was obtained by removing the distal 1-2 mm of the outer digit on the hind foot; samples were preserved in 96% ethanol and stored at −20℃ until the DNA extraction. The protocol was approved by the Ethics Committee of Saint-Petersburg State University (#131-03-9). All mole voles were released back to their exact point of capture.

Northern mole voles are highly social rodents. They live in extended family groups that can include up to 20-25 individuals, most of which are philopatric offspring of a few breeders (Evdokimov 2001). Both the presence of siblings in samples and the indiscriminate removal of putative siblings can affect the results of population genetic analyses (Anderson and Dunham 2008; Rodrıguez-Ramilo and Wang 2012; Parreira and Chikhi 2015; Waples and Anderson 2017). We resolved a tradeoff between two extreme options by sampling 1-7 individuals per putative burrow system (Table 1 and Table S1). Where possible, adults with the widest incisors were selected to limit the number of full siblings in our samples. The sampling strategies in Novosibirsk and Saratov were identical, which allowed for direct comparisons between these populations.

**Table 1.** Number of animals and corresponding family groups of *Ellobius talpinus* included in the analyses based on microsatellites and D-loop data.

| Population | Site | Microsatellites |  |  | D-loop |  |  | Total |
| --- | --- | --- | --- | --- | --- | --- | --- | --- |
|  |  | N<br>indivi<br>duals | N<br>grou<br>ps | Average N<br>individuals<br>per group<br>(± SD) | N<br>individ<br>uals | N<br>gro<br>ups | Average N<br>individuals<br>per group<br>(± SD) | N<br>individuals |
| Novosibirsk | NB | 33 | 13 | 2.5±1.5 | 30 | 15 | 2.0±1.0 | 43 |
|  | NI | 10 | 4 | 2.5±1.2 | 12 | 4 | 3.0±1.1 | 13 |
|  | **total** | **43** | **17** | **2.5±1.4** | **42** | **18** | **2.3±1.1** | **56** |
| Saratov | SR | 44 | 19 | 2.3±2.0 | 52 | 20 | 2.6±2.0 | 53 |
|  | SG | 9 | 5 | 1.5±0.8 | 9 | 5 | 1.5±0.8 | 9 |
|  | SF | 4 | 3 | 1.3±0.6 | 4 | 3 | 1.3±0.6 | 4 |
|  | **total** | **57** | **27** | **2.1±2.0** | **65** | **28** | **2.3±1.8** | **66** |

### 2.2. Primer selection and design

Initially, we attempted amplification of eleven microsatellite loci (Table 2): four were previously designed for Arvicolinae (Walser and Heckel 2008; Ishibashi et al. 1999), and seven loci were modified for *E. talpinus* in this research. New microsatellite loci were found by aligning published sequences from other voles (Ruda et al. 2009; Yue et al. 2009; Rikalainen et al. 2008) against the genome of *E. talpinus* (GenBank: LOJH00000000.1; Mulugeta et al. 2016) using BLAST algorithms. The found sequences were analyzed for their length and number of repeats in Geneious Prime 2021.0.1. We also used this software to find primers on the selected region of the locus. Next, the selected primers were verified using the OligoAnalyzerTool © 2021 (Integrated DNA Technologies, Inc.) to analyze possible hairpins and dimers formed. The primer pair specificity was tested by PCR amplification using *E. talpinu*s genomic DNA as a template. The length of the obtained fragments was estimated by electrophoresis in 1% agarose gel using ethidium bromide as a DNA stain. To test the specificity of the primers, the fragments were sequenced using the Sanger method.

**Table 2.**
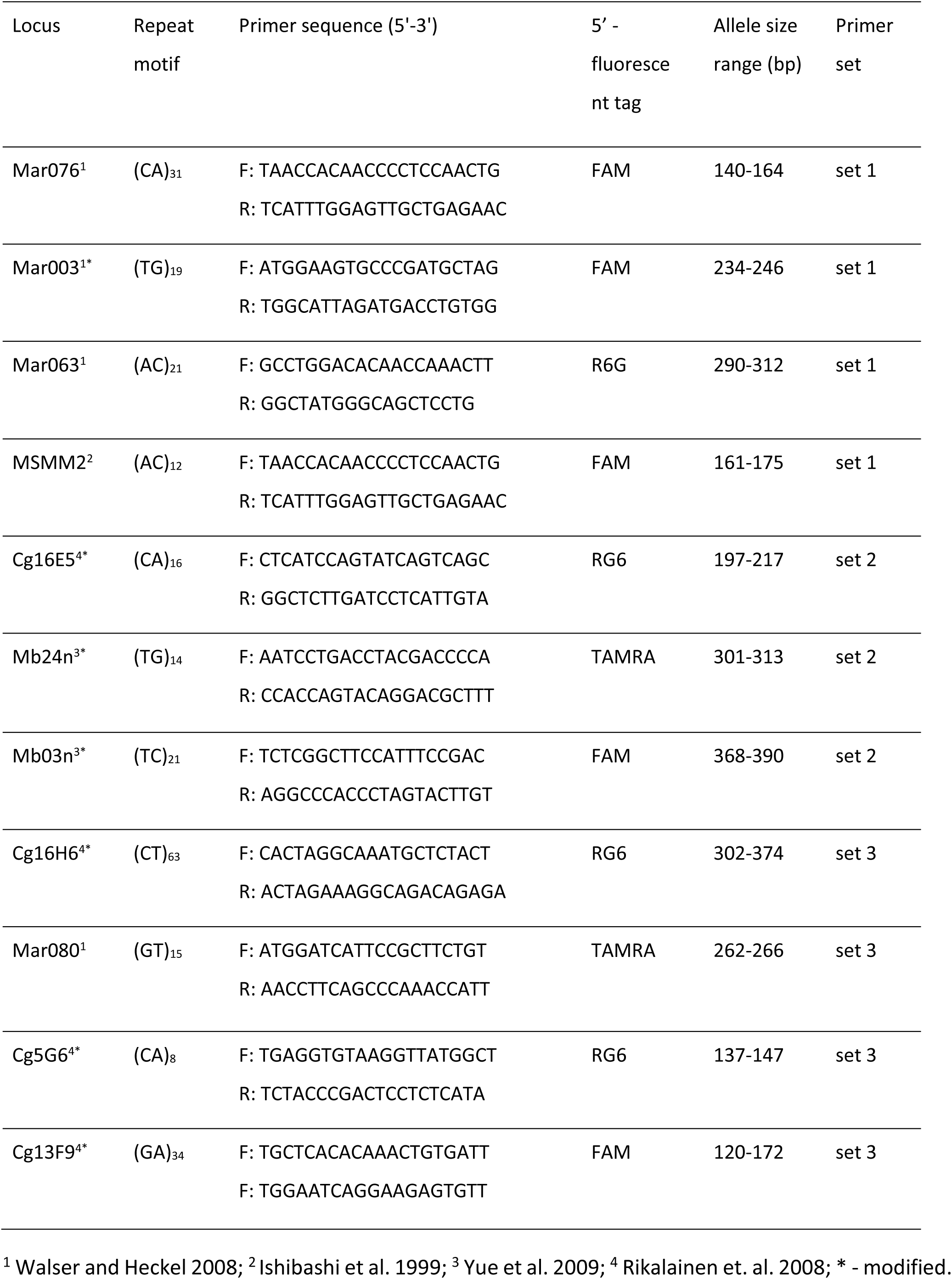
Primer sequences for microsatellite loci used in the study.

The amplification of the first variable fragment of the mitochondrial D-loop was performed with primers Ell3F (5’-TCAAGAAGGAAGGACCTACCС-3’) and Ell1R (5’-GGCTGATTAGACCCGTACCA-3’ which were designed using mitogenome of *E. talpinus* (NCBI: NC_054160.1) (Kuprina et al. 2024).

### 2.3. Molecular analysis

Total DNA extraction from phalanges followed a phenol–chloroform procedure, via digestion with proteinase K and ethanol precipitation (Sambrook et al. 1989). Some genomic DNA was extracted using spin-column from LumiPure Genomic DNA Extraction Kit (Lumiprobe, USA) following the manufacturer’s protocol.

Genotyping was performed at eleven microsatellite loci for a total of 100 samples (Table 3). The repeated (at least twice) genotyping of 20% of individuals yielded an error rate of 3%. Microsatellite markers were amplified in three multiplex sets (Table 2). Multiplex PCR was performed in 10 µl reaction volume containing 5 µl 2x QIAGEN Multiplex PCR Master Mix (QIAGEN, Germany), 2 µl 5x Q-Solution (QIAGEN), 1 µl primer mix (2 µM each primer), 1 µl genomic DNA (∼ 20 ng), and 1 µl RNase-free-water. Amplification started with an initial activation step at 95 °C for 15 min, followed by 35 cycles with denaturation at 94 °C for 30 s, annealing at 57 °C for 90 s, extension at 72 °C for 60 s and final extension at 60 °C for 30 min. Thermal cyclers T-100 or MJ Mini (Bio-Rad Laboratories, USA) were used. Fragment analysis was performed using the S550-LIS (COrDIS) size standard, according to the manufacturer’s recommendations.

Amplification of the mitochondrial D-loop fragment was performed in a 20 μl reaction mixture containing 1X buffer, 1.5 mM MgCl2 (both from Axon Labortechnik, Germany), 4 μM dNTPs, 0.2 μM of each primer, 1 U of HotStart Taq (Axon Labortechnik, Germany), and 60 ng DNA (2 μl). Amplification was conducted using a thermocycler MJ Mini (Bio-Rad Laboratories, USA) with the following program: (i) 13 min at 95 ◦C; (ii) 35 cycles of amplification consisting of 45 s at 95 ◦C, 30 s at 55 ◦C and 30 s at 72 ◦C; (iii) final extension of 3 min at 72 ◦C.

Sequencing and fragment analysis of the PCR products were performed using an ABI 3500 XL Genetic Analyser (Thermo Fisher Scientific, USA). Given the presence of multiple nuclear pseudogenes of the mitochondrial control region shown for mole voles (Kuprina et al. 2024), we paid special attention to the quality of the chromatograms: all samples were sequenced from both primers. Sequences whose haplotypes differed by 1-2 nucleotides and samples with unique haplotypes were analyzed repeatedly.

### 2.4. Data analyses

#### 2.4.1. Genetic diversity

The length of microsatellite fragments was estimated using GeneMarker 1.85 Demo (SoftGenetics) software. All gained peak values were rounded using Tandem 1.09 software (Matschiner et al. 2009). The locus Mar080 showed two alleles which differed by a single nucleotide, making the determination of allele length unreliable. The locus Cg13F9 exhibited strong stutters. Both loci were discarded due to these issues.

Micro-Checker 2.2.3 (Van Oosterhout et al. 2004) was used to detect scoring errors due to stuttering patterns, large-allele dropouts, and null alleles for the remaining loci. Calculations of allele frequencies, analyses of linkage disequilibrium, and tests of departure from Hardy-Weinberg equilibrium (HWE) were completed using GENEPOP 4.7.5 (https://genepop.curtin.edu.au/; Raymond and Rousset 1995). We used a Markov chain method, with 10,000 dememorization steps, 500 batches, and 5,000 iterations per batch. Holm’s sequential Bonferroni *p*-value correction for multiple tests was carried out. These analyses were performed separately for each of the two populations and for each of the two sites with sample size greater than 10 (NB and SR).

Based on the loci suitable for population analysis, estimates of observed (*Ho*) and unbiased expected heterozygosity (*uHe*), and inbreeding coefficients (*Fis*) were obtained for each population and each site in GenAlex 6.5 (Peakall and Smouse 2012). To estimate allele richness while accounting for uneven sample sizes (Ar), a rarefaction analysis was conducted using HP-rare 1.0. (Kalinowski, 2005). We used the Wilcoxon Signed Rank Test (STATISTICA 12) to compare rarefied *Ar*, *Ho* and *uHe* between the Novosibirsk and Saratov populations and between the NB and SR sites (which are comparable in trapping areas and sample sizes).

The chromatograms of the mtDNA D -loop were manually checked with Chromas 2.6.6, and the sequences were aligned in MEGA 11 (Tamura et al. 2021). Genetic diversity, number of haplotypes, polymorphic sites, haplotype (*Hd*), and nucleotide diversity (*π*) indices were determined for each sampling site using DnaSP v5.10 (Librado and Roxas 2009).

To visualize haplotype variability, a haplotype network was obtained from the TCS analysis (Clement et al. 2002) using the program PopART 1.7 (Leigh and Bryant 2015). Phylogenetic relationships among D-loop haplotypes of *E. talpinus* were estimated under Bayesian inference (BI) using MRBAYES v3.2 (Ronquist et al. 2012) (10,000,000 generations, 25% burn-in) and visualized in FigTree v.1.4.3 (tree.bio.ed.ac.uk/software/figtree/). In addition to the haplotypes obtained in this study, three sequences from GenBank were included: the *E. talpinus* D-loop fragment (individual from the Rostov Region, NCBI: NC_054160), *E. talpinus* pseudogene (individual from the Novosibirsk Region, NCBI: OR662054), and *E. tancrei* as an outgroup (Southwestern Tajikistan, NCBI: OR662053). The best-fit substitution model (HKY + G) was determined using MEGA 11.

#### 2.4.2. Genetic structure and gene flow

We examined population subdivision by conducting the analysis of molecular variance (AMOVA; Excoffier et al. 1992). This analysis involved partitioning genetic variation hierarchically between two populations, among sites within populations, and among individuals within sites. Among several measures proposed for description of microsatellite-based genetic structure, Fst (Wright 1965) is the most commonly used. It is particularly effective in assessing the low degree of differentiation that might be expected at a small spatial scale, such as within the Saratov population. However, Fst tends to underestimate the differentiation of highly divergent populations, such as Novosibirsk and Saratov in this study. In these situations, Rst (Slatkin 1995) is more appropriate (Balloux and Lugon-Moulin 2002). Therefore, we used both Rst- and Fst-based AMOVAs implemented in GenAlex 6.5 (Peakall and Smouse 2012). For haplotype data, AMOVA was conducted in Arlequin v.3.5.2.2 (Excoffier and Lischer 2010) with 10,000 permutations. We assessed genetic differentiation adjusting p-values with the Bonferroni correction. In addition, we explored the genetic structure using discriminant analysis of principal components (DAPC; Jombart et al. 2010; Jombart and Collins 2015). This multivariate clustering method is free of the assumptions of population genetics, particularly Hardy-Weinberg equilibrium, and was therefore preferred over approaches that rely on algorithms based on predefined population genetics models, such as STRUCTURE (Pritchard et al. 2003) and BAP (Corander et al. 2003). DAPC analyses were conducted twice using the R package “adegenet 2.0.0” (Jombart and Collins 2015) to assess the influence of a priori groupings on the results. In the initial analysis, sites were used as a priori groups, and the “find.clusters” function was applied to determine the number of clusters (*K*) *de novo*, with the optimal K associated with the lowest Bayesian Information Criterion (BIC) value. Additionally, we conducted analyses without predefined groups and used various parameterizations to test stability of the results (Meirmans 2015; Thia 2022). Specifically, grouping of individuals was inferred from PCA including 30, 50, and 80 principal components (PCs), and the optimal number of PCs retained in each DAPC was set to K-1 (Thia 2022). The resulting clusters were visualized as DAPC scatterplots. All R packages used in this study were executed in RStudio IDE 2023.6.1+524 (2020) and R 4.4.1 (R Core Team 2023).

To further explore the fine-scale genetic structure, we tested whether patterns of isolation by distance were present within the largest sites within each population (NB in Novosibirsk and SR in Saratov). First, we used the Mantel test to estimate the correlations between genetic and geographic distances (log-transformed data). Microsatellite data were analyzed using GenAlEx 6.5 (Peakall and Smouse 2012), while haplotype data were processed in R with the “mantel” function, employing Pearson’s correlation and 9999 permutations. Second, we conducted spatial autocorrelation analyses in GenAlEx 6.5 to estimate the distances at which relatedness (based on the microsatellite data) is positive. We used distance classes of 50, 100, 300, 600, and 1000 m, setting the first class slightly larger than the estimated home range size. Statistical testing was based on 999 random permutations of individual genotypes.

#### 2.4.3. Historical changes in population size

We investigated aspects of population history and demography using both microsatellite and haplotype data. To eliminate the confounding effect of geographic structure, neutrality tests were first carried out individually for the Novosibirsk and Saratov populations, and then for the NB and SR sites within each population. Given that these tests have low statistical power when sample sizes of loci and/or individuals are small (Cornuet and Luikart 1996; Peery et al. 2012; Hoban et al. 2013), we focused on the population history of only two sites with the largest sample sizes.

Several microsatellite-based methods were used. First, we analyzed allele frequency distributions, as a recently bottlenecked population is expected to show a lower number of alleles in the low frequency class (< 0.1) compared to one or more intermediate frequency classes (Luikart et al. 1998). Second, we performed the M-ratio test (Garza and Williamson 2001) which is based on the assumption that a bottleneck event will result in loss of alleles from allele size categories. It allows the detection of bottleneck events that took place 5-100 generations ago. Estimates of M-ratio were calculated in Arlequin v3.5.2.2 (Excoffier and Lischer 2010) and compared to Mc = 0.68 (the critical value for the dataset based on seven loci or more: - Garza and Williamson 2001). The third method relies on assessing an excess of heterozygosity compared to the expected equilibrium values for a given allelic diversity (Cornuet and Luikart 1996). A one-tailed Wilcoxon test implemented in BOTTLENECK v1.2.02 (Cornuet and Luikart 1996) was used to determine if a statistically significant number of loci displayed excess of heterozygosity compared to expectations (number of iterations = 3000, variance in TPM = 10). We repeated this analysis with a range of values for the proportion of multi-step mutations (pg), from the strict SSM model to pg = 50% (Peery et al. 2012).

We also tested the hypothesis of demographic stability of the populations with several statistics appropriate for sequence data. First, we used Tajima’s D (Tajima 1989) and Fu’s Fs (Fu 1997) statistics using DnaSP v. 5.10 (Librado and Rozas 2009) to see whether *E. talpinus* data conformed to expectations of neutrality or departed from neutrality due to factors such as a population bottleneck or expansion. The significance of these tests was assessed by comparing the observed values with their empirical distributions, generated from 1000 replicate coalescent simulations in the ‘coalescent simulations’ module in DnaSP, where the number of segregating sites held constant under a constant population size hypothesis. In a constant-size population, the expected values of these statistics are nearly zero; significant negative values suggest a sudden population expansion, whereas significant positive values indicate processes such as a population subdivision or recent population bottlenecks. Second, we quantified the smoothness of the observed mismatch distributions using the raggedness index according to (i) the demographic expansion model (DEM) and (ii) the spatial expansion model with population subdivision (SEM) implemented in Arlequin 3.5 (Excoffier and Lischer 2015). Small raggedness values indicate a population that has experienced sudden expansion whereas higher values suggest stationary or bottlenecked populations. Third, we tested the goodness-of-fit of the observed mismatch distributions to those expected under (i) the demographic expansion model (DEM) and (ii) the spatial expansion model with population subdivision (SEM) implemented in Arlequin 3.5. A significance was assessed by comparing the observed raggedness and sum of squared deviations (SSD) with a distribution of values drawn from data simulated using parameters optimized from the observed data under each demographic scenario (Slatkin and Hudson 1991; Rogers and Harpenting 1992).

## 3. RESULTS

We successfully genotyped 100 samples using microsatellites and identified D-loop haplotypes in 107 individuals from five sites (Table 4). Two microsatellite markers (Mar080 and Cg13F9) did not amplify well in individuals from the Novosibirsk population, so only nine markers were used in the analyses.

**Table 4.** Genetic variation in populations of *Ellobius talpinus* from the Novosibirsk (N, sites NB and NI) and Saratov populations (sites SR, SG, and SF). *A (±SE)* - number of alleles per locus, *Ar* - rarefied allelic richness (min. number of alleles = 8, based on the smallest population sample size for SF), *Ho (±SE)* - observed heterozygosity, *uHe (±SE)* - unbiased expected heterozygosity; *Fis* - inbreeding coefficients; H – number of haplotypes; *Hd (±SD)* – haplotype diversity; *S* (%) – number of polymorphic sites; *π (±SD)* – nucleotide diversity; *k (±SD)* - average number of nucleotide differences

| Site/<br>population | Microsatellites |  |  |  |  | D-loop |  |  |  |  |
| --- | --- | --- | --- | --- | --- | --- | --- | --- | --- | --- |
| | *A* | *Ar* | *Ho* | *uHe* | *Fis* | *H* | *Hd* | *S* | $\pi$ | *k* |
| NB | 4.11<br>(0.841) | 2.7<br>7 | 0.337<br>(0.059) | 0.506<br>(0.094) | 0.300<br>(0.062) | 3 | 0.191<br>(0.093) | 2<br>(0.5) | 0.0005<br>(0.0003) | 0.20<br>(0.06) |
| NI | 3.56<br>(0.729) | 3.0<br>3 | 0.358<br>(0.096) | 0.522<br>(0.107) | 0.327<br>(0.090) | 1 | 0 | 0 | 0 | 0 |
| Novosibirsk,<br>total | 5.00<br>(1.160) | 2.9<br>0 | 0.339<br>(0.064) | 0.550<br>(0.098) | 0.351<br>(0.069) | 3 | 0.138<br>(0.071) | 2<br>(0.5) | 0.0004<br>(0.0002) | 0.14<br>(0.04) |
| SR | 8.11<br>(1.218) | 4.0<br>5 | 0.662<br>(0.049) | 0.720<br>(0.047) | 0.082<br>(0.026) | 7 | 0.789<br>(0.030) | 31<br>(8.1) | 0.033<br>(0.0007) | 12.7<br>(34.1) |
| SG | 3.44<br>(0.338) | 2.8<br>4 | 0.556<br>(0.072) | 0.587<br>(0.040) | 0.020<br>(0.137) | 1 | 0 | 0 | 0 | 0 |
| SF | 3.44<br>(0.444) | 3.4<br>4 | 0.583<br>(0.110) | 0.635<br>(0.100) | 0.086<br>(0.115) | 2 | 0.500<br>(0.265) | 17<br>(4.4) | 0.022<br>(0.0155) | 8.50<br>(24.9) |
| Saratov, total | 8.89<br>(1.500) | 3.4<br>4 | 0.640<br>(0.051) | 0.760<br>(0.038) | 0.165<br>(0.043) | 9 | 0.834<br>(0.020) | 34<br>(8.8) | 0.034<br>(0.0008) | 13.0<br>(35.2) |

### 3.1. Genetic diversity

The linkage disequilibrium test revealed significant linkage for three pairs of loci in Novosibirsk (Mar076 - Mar003, Cg16H6 - Mar003, and Mar076 - MSMM2) and for three pairs of loci in Saratov (Mar063 - MSMM2; Cg16H6 - Mb03n, and Cg16H6 - MSMM2). Analysis at a smaller scale found significant linkage disequilibrium for only one pair of loci (Mar076 - MSMM2) at NB. All observed deviations were population-specific, suggesting that the disequilibrium was not due to physical linkage, but rather to substructure or the presence of close relatives in the sample. Therefore, none of the loci were dismissed.

Micro-Checker 2.2.3 did not detect any genotyping errors but warned of possible presence of null alleles at loci Mar063, Mar003, Cg5G6, and Mb03n in Saratov, and at all but two loci (Cg5G6 and Cg16E5) in Novosibirsk (Table S2). The presence of null alleles, especially at a high frequency, as observed in Novosibirsk population, should result in the occurrence of blank genotypes (putative null homozygotes). In Novosibirsk, possible null alleles were detected at only two loci in two samples. However, these samples were not amplified for some other loci where null alleles were not detected by the program, suggesting that this may be due to the low quality of the DNA samples. In Saratov, all samples amplified successfully at the loci with putative null alleles. Combined with the population-specificity of most loci with identified null alleles and the almost complete lack of locus-specificity in Novosibirsk, these findings allowed us to reject the null allele hypothesis. Several factors could lead to t false detection of null alleles, including generations sampled, sample size, the level and duration of bottlenecks, kin clustering, and non-random mating (Dąbrowski et al. 2014). Therefore, we retained all nine loci for further analyses.

Observed heterozygosity was lower than expected at all sites (Table 4). We found a significant deviation from HWE at all polymorphic loci but Cg16E5 in the Novosibirsk population and at four loci (Mar003, Mb03n, Cg5G6 and Mb03n) in the Saratov population (Table S2). On a smaller scale, significant deviations from HWE were found at six loci (Mar076, Mar063, Cg16H6, Mb03n, Mb24n, and MSMM2) in Novosibirsk and at two loci (Mar003 and Mb03n) in Saratov (Table S2). The Fis values were significantly higher in Saratov compared to Novosibirsk (Z = 2.38; n = 8; *p* = 0.017) and higher at SR than at NB (Z = 2.52; n = 8; *p* = 0.012).

A total of 94 alleles were detected across nine microsatellite loci. All loci were polymorphic in Saratov, while the Novosibirsk population had one monomorphic locus (Cg5G6). There was a clear genetic distinction between the two populations with unique alleles observed at all loci. The Novosibirsk population contained 13 unique alleles of six loci, whereas the Saratov population had 43 unique alleles of nine loci. Genetic diversity parameters for each population and sites are shown in Table 4 and Table S2. The rarefied allelic richness tended to be higher in Saratov than in the Novosibirsk population (Z=1.84, n = 9, *p* = 0.066). Both observed and unbiased expected heterozygosity were significantly higher in Saratov (*Ho*: Z=2.67, n = 9, *p* = 0.008; *uH*e: Z = 2.31; n = 9; *p* = 0.021). The comparison between two sites with similar area and sample sizes (NB and SR) yielded a similar pattern, with significantly higher diversity indexes at SR than at NB (*Ar*: Z = 2.67, n = 9; *p* = 0.008; *Ho*: Z = 2.67, n = 8, *p* = 0.008; *uHe*: Z = 2.55; n = 9, *p* = 0.011). There was no tendency for allele lengths in Saratov to be greater than those in Novosibirsk (Wilcoxon signed-ranks test, T = 22, n = 9, *p* = 0.953), suggesting that the observed differences in variability were not due to consistent biases in microsatellite size.

A 385 bp fragment of the mitochondrial D-loop region was sequenced for 107 individuals. In total, 46 polymorphic sites, including one indel, were identified defining 12 different haplotypes, nine at Saratov and three at Novosibirsk (Table S3). No haplotype was present in both populations. Overall haplotype diversity was 0.81 (SD = ±0.03), and the nucleotide diversity was 0.042 (SD = ±0.001), indicating a high level of polymorphism. All diversity indexes were higher in Saratov than in the Novosibirsk population, and higher at SR than at NB (Table 4).

The topology of the main clades of the Bayesian phylogenetic tree including the detected haplotypes, an additional *E. talpinus* haplotype from GenBank, the pseudogene of *E. talpinus* and D-loop fragment of *E. tancrei*, was as expected (Fig. 3). D-loop pseudogene was basal relative to all other *E. talpinus* haplotypes. These last were grouped into three divergent lineages corresponding to geographic populations (Novosibirsk, Saratov, and Rostov regions), though the relationships between these lineages were poorly resolved. Within the Saratov population, four well-differentiated clades, each including frequently occurring haplotypes, were revealed by both Bayesian phylogenetic analysis (Fig. 3) and TCS network analysis (Fig. 4). In contrast, the Novosibirsk population was nearly fixed for a single haplotype, found in 93% of individuals (Fig. 4).

**Figure 3.**
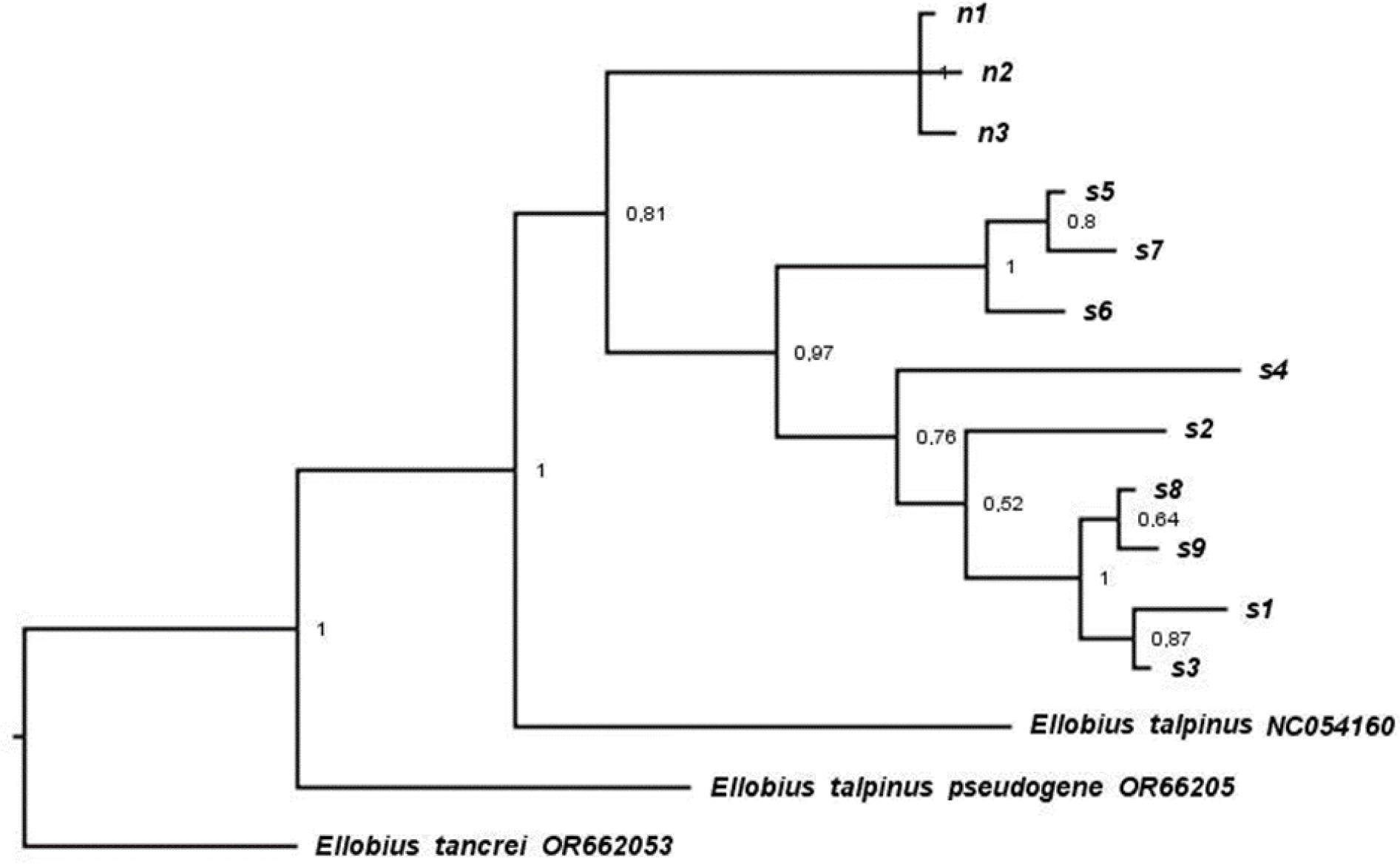
Bayesian tree of *Ellobius* haplotypes based on a 385 bp fragment of mtDNA. Twelve D-loop haplotypes of *E. talpinus* (*s1 - s9*; *n1 - n3*) were detected in this study, and three additional haplotypes from Genbank: D-loop fragments of *E. tancrei* and *E. talpinus*, and one D-loop pseudogene of *E. talpinus*. Values near nodes indicate Bayesian posterior probabilities.

**Figure 4.**
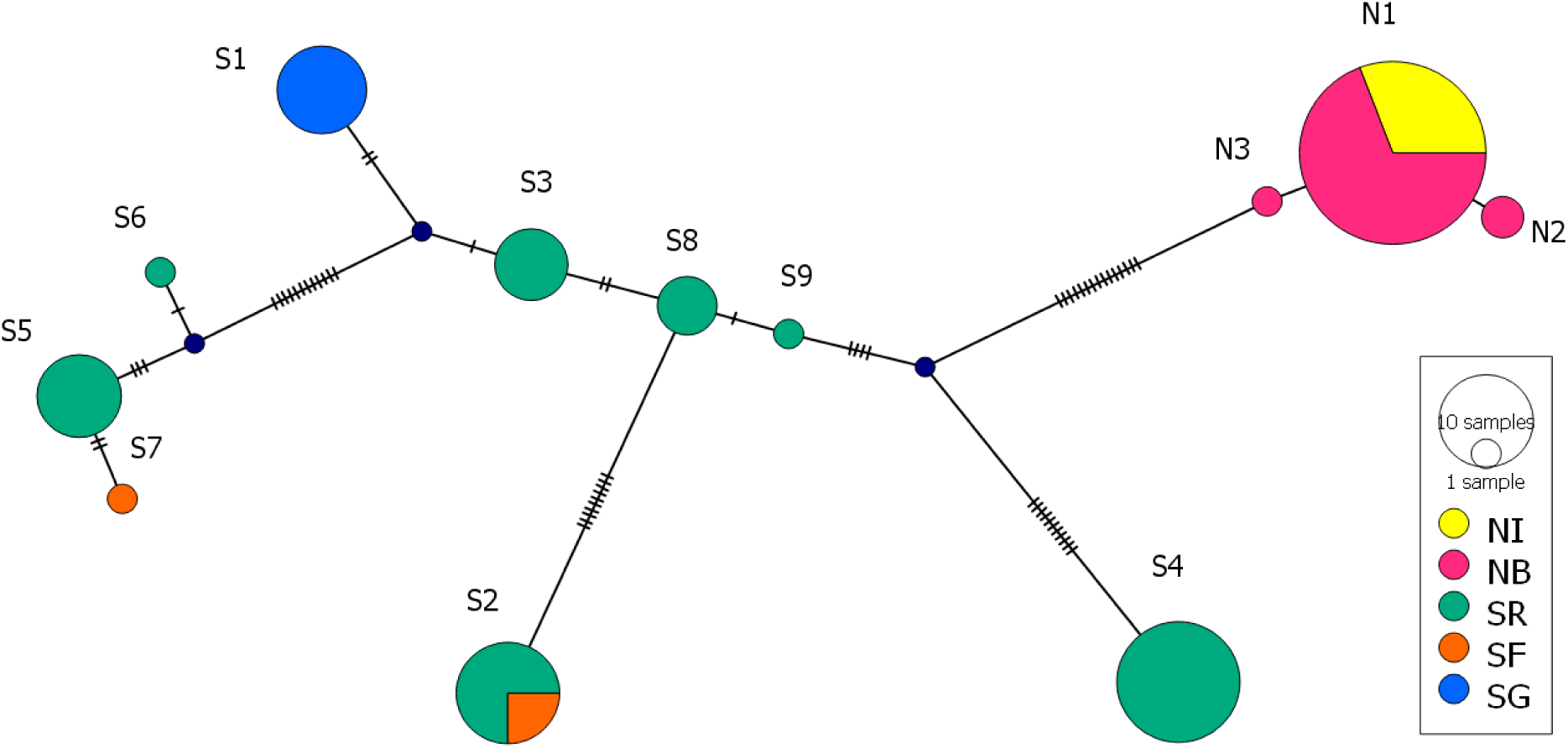
TCS network of D-loop *Ellobius talpinus* haplotypes. Hatch marks along the lines indicate the number of mutations between haplotypes. The black circles between haplotypes correspond to intermediate haplotypes as calculated by the program.

### 3.2. Population structure and gene flow

According to the results of both microsatellite- and D-loop-based AMOVA, the majority of the variance was observed within sites, rather than among populations or sites (Table 6). Based on pairwise Fst for microsatellite data, all site pairs within each population were significantly differentiated (Table 7). The Rst-based AMOVA for microsatellites caught significant differentiation only between NI and NB (Table 7). The D-loop-based AMOVA сonfirmed the isolation of SG from the other two sites within the Saratov population but failed to reveal the significant differentiation between SF and SR (Table 7). No structure was found within the Novosibirsk population, probably due to the very low level of polymorphism (Table 7).

**Table 5.**
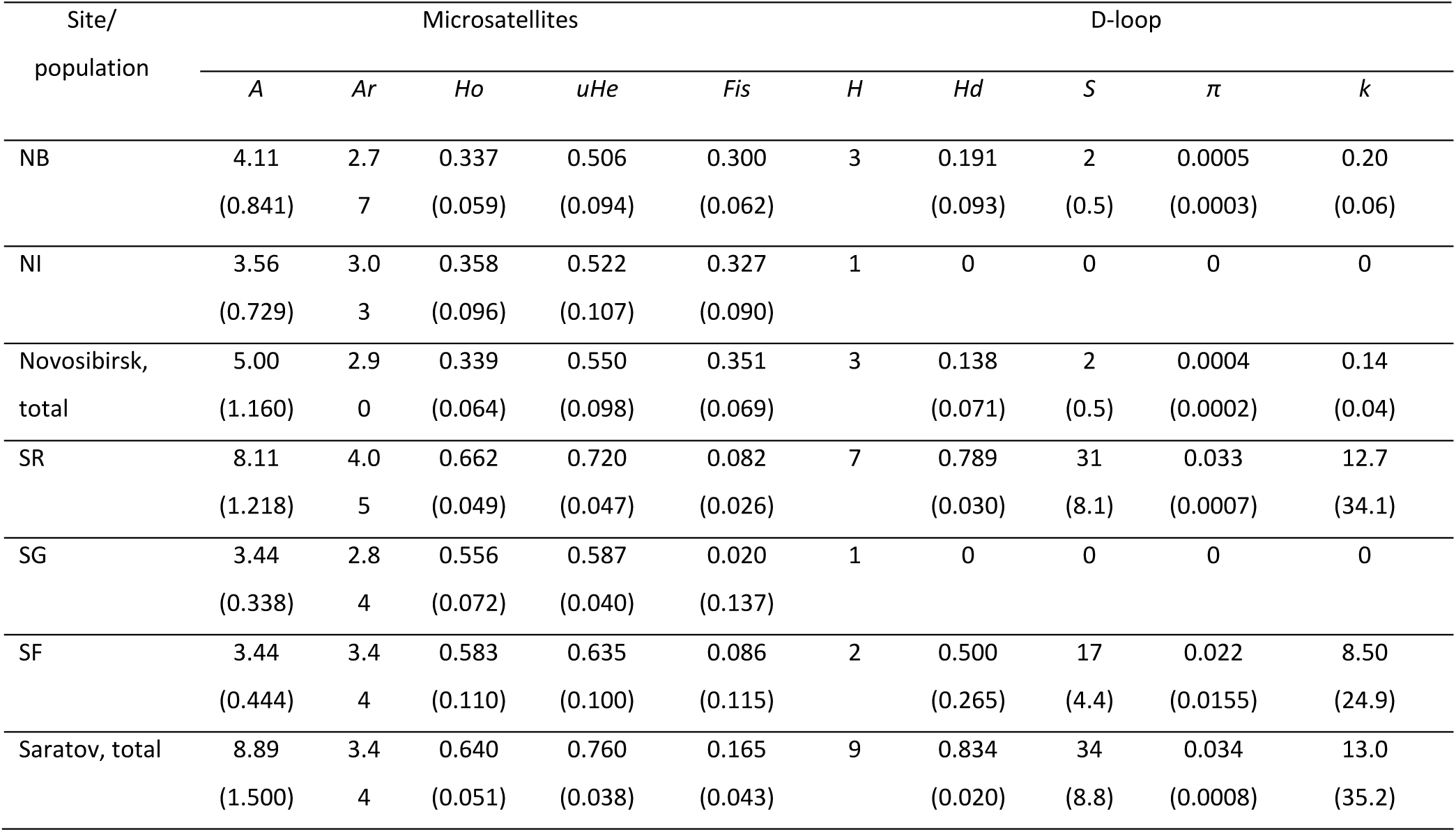
Genetic variation in populations of *Ellobius talpinus* from the Novosibirsk (N, sites NB and NI) and Saratov populations (sites SR, SG, and SF). *A (±SE)* - number of alleles per locus, *Ar* - rarefied allelic richness (min. number of alleles = 8, based on the smallest population sample size for SF), *Ho (±SE)* - observed heterozygosity, *uHe (±SE)* - unbiased expected heterozygosity; *Fis* - inbreeding coefficients; H – number of haplotypes; *Hd (±SD)* – haplotype diversity; *S* (%) – number of polymorphic sites; *π (±SD)* – nucleotide diversity; *k (±SD)* - average number of nucleotide differences

**Table 6.** Results of the Analysis of Molecular Variance (AMOVA) of microsatellites and mtDNA D-loop sequences in *Ellobius talpinus* from two populations and five sites. DF - degrees of freedom, SS - sum of squares, Var - estimated variance, % - percentage of molecular variance.

| Source of Variation | DF | SS | Var | % | Fixation indices | p-value |
| --- | --- | --- | --- | --- | --- | --- |
| Among populations |  |  |  |  |  |  |
| microsatellites, Fst | 1 | 93.54 | 0.490 | 12 | 0.122 | 0.001 |
| Rst | 1 | 14594.1 | 128.5 | 29 | 0.289 | 0.002 |
| D-loop Fst | 1 | 13.51 | 0.174 | 34 | 0.327 | 0.097 |
| Among sites within populations |  |  |  |  |  |  |
| microsatellites, Fst | 3 | 58.81 | 0.685 | 17 | 0.194 | 0.001 |
| Rst | 3 | 2919.5 | 26.9 | 6 | 0.089 | 0.001 |
| D-loop Fst | 3 | 5.88 | 0.133 | 24 | 0.576 | 0.001 |
| Within sites |  |  |  |  |  |  |
| microsatellites, Fst | 95 | 554.50 | 2.844 | 71 | 0.293 | 0.001 |
| Rst | 95 | 29917.7 | 27.5 | 65 | 0.353 | 0.001 |
| D-loop Fst | 102 | 23.11 | 0.227 | 42 | 0.369 | 0.001 |

**Table 7.** Pairwise genetic differentiation estimates in *Ellobius talpinus* across two populations and five sites. Significant fixation indices, after Bonferroni correction, are shown in bold.

| Pair of sites | Differentiation estimate | Fixation index | p-value |
| --- | --- | --- | --- |
| NB-NI | Microsatellite Fst | **0.188** | **0.001** |
|  | Microsatellite Rst | **0.176** | **0.002** |
|  | D-loop Fst | -0.004 | 0.549 |
| SR-SF | Microsatellite Fst | **0.170** | **0.001** |
|  | Microsatellite Rst | 0.050 | 0.123 |
|  | D-loop Fst | 0.200 | 0.020 |
| SR-SG | Microsatellite Fst | **0.202** | **0.001** |
|  | Microsatellite Rst | 0.020 | 0.180 |
|  | D-loop Fst | **0.458** | **0.001** |
| SG-SF | Microsatellite Fst | **0.249** | **0.001** |
|  | Microsatellite Rst | 0.202 | 0.017 |
|  | D-loop Fst | **0.852** | **0.002** |

Using the five sites as *a priori* groups and four retained DA, the first PC accounted for 79% of the variance. It differentiated the Novosibirsk population from the Saratov population and NB from NI (Fig. 5A). The second PC accounted for 14% of the variance and distinguished the three Saratov sites from one another (Fig. 5A). All genotypes were correctly assigned to their groups. Without predefined groups, the optimal K varied from six to eight (see Fig. S1 for BIC plots). Respectively, we conducted DAPC with the number of retained PCs (*K*-1) five, six and seven. The first PC accounted for 65% (*K* = 8) to 69% (*K* = 6) of the variance and clearly differentiated the Novosibirsk population from the Saratov population (Fig. 5B). The second PC accounted for 16-16.6% of the variance and differentiated groups within each population (Fig. 5B). Within the Novosibirsk population, one cluster consistently contained eight animals from NI, while the second cluster included the remaining two individuals from NI along with some of the NB individuals. Within the Saratov population, the first cluster consistently included all nine individuals from SG; the second cluster contained four animals from SF and some of the SR individuals. The membership of the other clusters was inconsistent and did not display any particular spatial pattern in either SR or NB. Taken together, these results revealed differentiation between NI and NB, the isolation of SG from the other two Saratov sites, and the lack of spatial structure within the NB site and within the area covering SR and SF.

**Figure 5.**
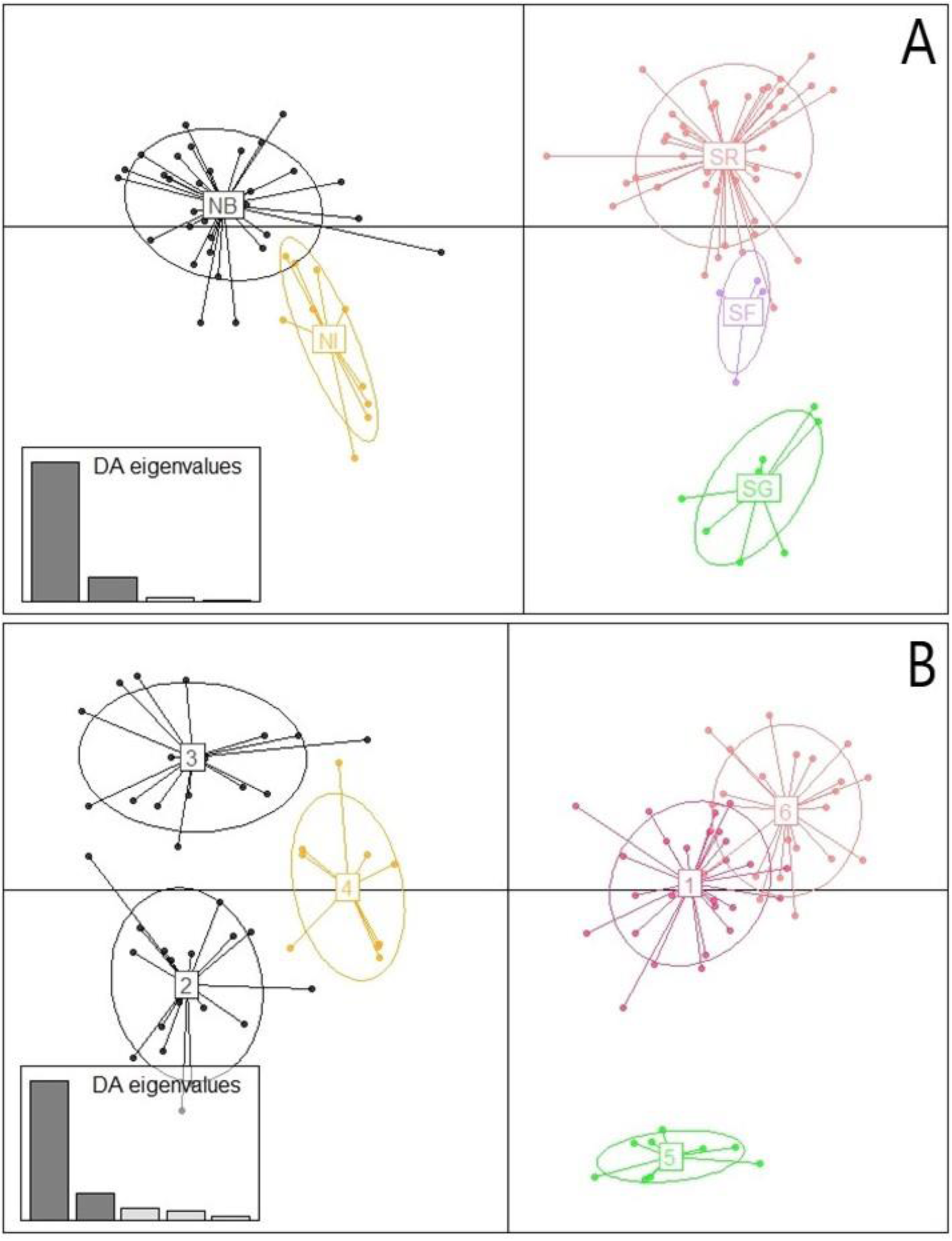
Scatterplot of the DAPC results for 100 *Ellobius talpinus* (A) with five sites as prior clusters (sampling site names are abbreviated as in Table 1) and (B) with no a priori population information (one of the obtained configurations). Dots represent different individuals. The size of the inertia ellipses encompasses 67% of individuals (default value). In (B), sites are assigned to clusters as follows: Cluster 1: 22 individuals from SR and 4 from SF; Cluster 2: 18 individuals from NB; Cluster 4: 9 individuals from NI; Cluster 3: 15 individuals from NB and 1 from NI; Cluster 6: 22 individuals from SR; Cluster 5: 9 individuals from SG.

**Figure 6.**
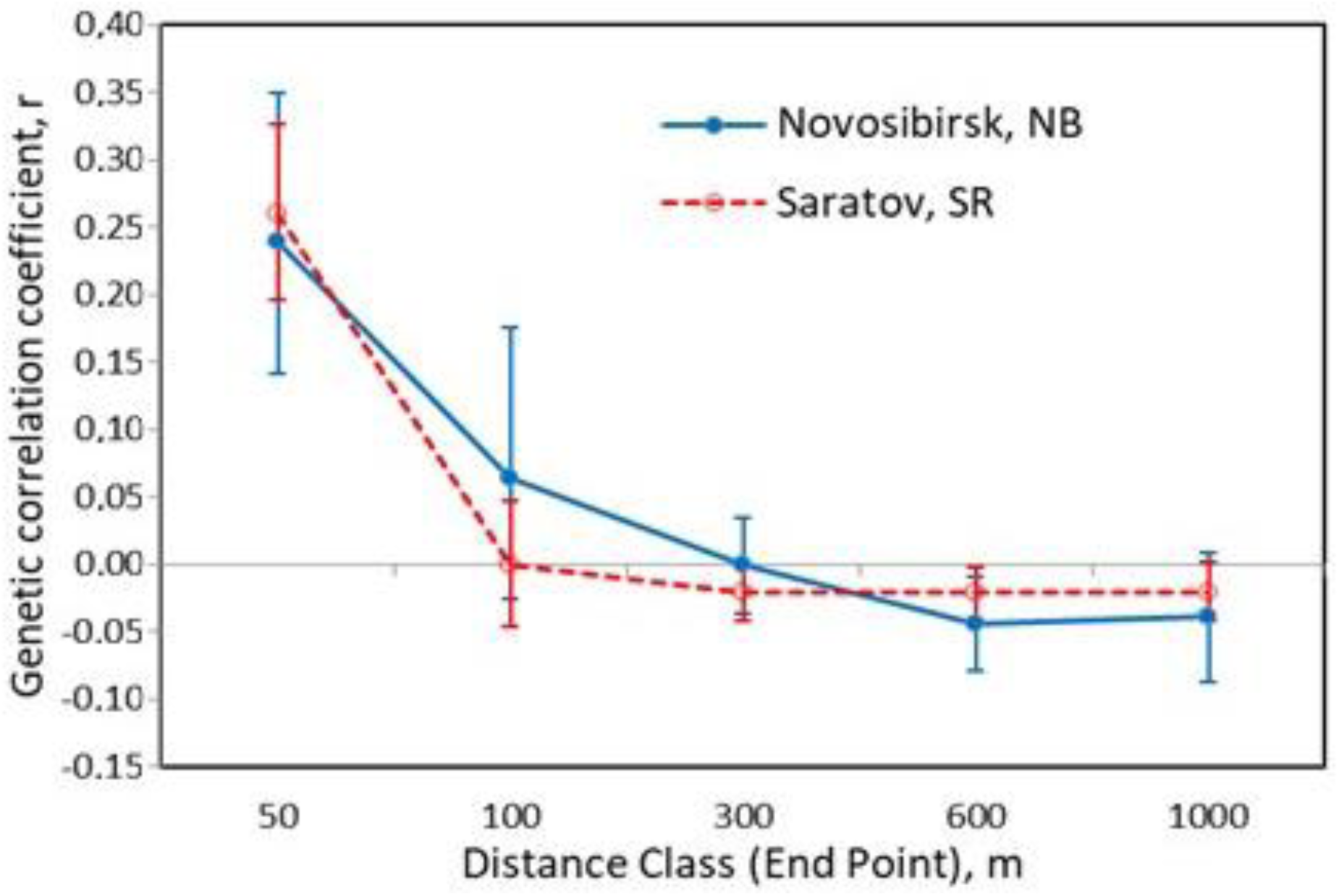
Correlograms showing genetic correlation (r) as a function of spatial distance for *Ellobius talpinus* from two populations. The 95% confidence error bars about r were determined by bootstrapping.

The results of the D-loop-based and microsatellite-based Mantel tests were similar. The correlation between genetic and geographic distances was insignificant at site NB of the Novosibirsk population (microsatellites: r = 0.124; *p* = 0.076; D-loop: r = 0.054; *p* = 0.191). In contrast, a significant spatial genetic structure was detected at site SR of the Saratov population (microsatellites: r = 0.183; *p* = 0.005; D-loop: r = 0.110; *p* = 0.020). The correlogram (Fig. 7) suggested that the lack of significant correlation at NB is most likely due to the high variability of the level of relatedness between individuals at short distances (up to 100 m). In both the Novosibirsk and Saratov populations, significantly positive r-values were found only within the 50 m distance class, likely reflecting family structure. In Saratov, r- values showed little change at distances greater than 100 m, while in Novosibirsk the average values continued to gradually decrease up to 600 m (Fig. 7).

### 3.3 Historical changes in population size

In both populations and at both large sites, alleles in the lowest frequency class were more abundant than alleles in any intermediate frequency classes (Fig. S2). In the Saratov population and at the SR site, the distribution was clearly L-shaped, with 63% and 58% of alleles, respectively, occurring at frequencies of < 0.10. In the Novosibirsk population and the NB sites, the proportions of rare alleles were lower (44% and 37%, respectively), but the patterns did not conform to Luikart’s criteria (Luikart et al. 1998) for detecting bottlenecks.

For the Novosibirsk population, the BOTTLENECK software did not show signs of a recent bottleneck under any mutation model (Table 8). For the Saratov population, a significant heterozygote excess indicative of a recent bottleneck was found only under TPM with pg = 0.5 (Table 8). The M-test did not reveal signs of a bottleneck in either of the two populations or sites (Table 8).

**Table 8.** Results of the one-tailed Wilcoxon test for heterozygote excess and the Garza–Williamson index for two populations and two largest sites (NB and SR) of *Ellobius talpinus*. N - sample size; SMM - stepwise mutation model; TPM - two-phase mutation models with various proportions of multi-step mutations; M - Garza–Williamson index.

| Parameter | Novosibirsk,<br>total | NB | Saratov,<br>total | SR |
| --- | --- | --- | --- | --- |
| N | 43 | 33 | 57 | 44 |
| SMM | 0.63 | 0.37 | 0.82 | 0.85 |
| TPM-0.1 | 0.527 | 0.273 | 0.410 | 0.715 |
| TPM-0.3 | 0.421 | 0.156 | 0.248 | 0.410 |
| TPM-0.5 | 0.230 | 0.098 | 0.019 | 0.367 |
| M (±SD) | 0.82 ± 0.17 | 0.82 ± 0.20 | 0.88 ± 0.13 | 0.91 ± 0.12 |

Non-significant negative Tajima’s D and Fu’s Fs values were obtained for the Novosibirsk population and the NB site (Table 9). In this case, negative D appears to be caused by a strong bottleneck (Fay and Wu 1999). The mismatch distributions were highly left-skewed, reflecting the absolute predominance of a single haplotype and its weak divergence from additional haplotypes. According to the mismatch distribution analysis, both expansion models could not be rejected for either the pooled Novosibirsk sample or site NB (Table 9; Fig. S3 A for DEM).

**Table 9.** Sequence-based demographic statistics for two populations, two largest sites and two polymorphic haplogroups of *Ellobius talpinus*. r - Harpending’s Raggedness index; SSD - sum of squared deviations between observed and expected mismatch distribution. Asterisks indicate levels of significance: * ≤ 0.05, ** ≤ 0.01, *** ≤ 0.001.

| Population/site | N | Tajima's D | Fu's FS | Demographic expansion |  | Spatial expansion |  |
| --- | --- | --- | --- | --- | --- | --- | --- |
|  |  |  |  | *r* | SSD | *r* | SSD |
| Novosibirsk, total | 42 | -1.3 | -2.09 | 0.552 | 0.000 | 0.552 | 0.0003 |
| NB | 30 | -1.26 | -1.67 | 0.434 | 0.001 | 0.434 | 0.000 |
| Saratov, total | 65 | 2.48** | 13.7** | 0.107*** | 0.054*** | 0.107 | 0.037 |
| SR | 52 | 2.66** | 14.9** | 0.306*** | 0.142*** | 0.306 | 0.102* |
| Saratov,<br>haplogroup A<br>(s1+s3+s8+s9) | 21 | 1.56 | 2.60 | 0.264 | 0.081 | 0.264 | 0.052 |
| Saratov,<br>haplogroup B<br>(s5+s6+s7) | 14 | -1.96** | 0.54 | 0.596 | 0.058 | 0.596 | 0.018 |

In both the Saratov population and site SR, Tajima’s D and Fu’s Fs were strongly positive, which may indicate a recent bottleneck or population subdivision. The mismatch distribution statistics obtained for Saratov and SR were not consistent with DEM (Table 9). A highly ragged trimodal distribution (Fig. S3 B) as well as positive D potentially could be due to a mixture of several distinct lineages, the presence of which in the Saratov sample was evident from the TCS network and phylogenetic relationship analyses (Figs. 3 and 4). Such a mixture would violate the assumptions of coalescent theory if analyzed as one ‘genetic’ population (Alvarado-Bremer et al. 2005). Therefore, we additionally carried out mismatch analysis and neutrality tests separately on two polymorphic and well supported lineages: haplogroup A, which includes haplotypes *s1*, *s3*, *s8*, and *s9*, and haplogroup B, which includes *s5*, *s6*, and *s7* (Figs. 3 and 4). Tajima’s D was not significantly different from zero for haplogroup A and was significantly negative for haplogroup B. Fu’s values were negative but non-significant, whereas the mismatch distribution was consistent with both DEM and SEM for A and B haplogroups (Table 9).

## 4. DISCUSSION

### 4.1. Comparison of genetic diversity in the Novosibirsk and Saratov populations

To our knowledge, this is the first study to examine intra-population genetic diversity and genetic differentiation using highly polymorphic, selectively neutral markers in any *Ellobius* species. We investigated the genetic patterns in two *E. talpinus* populations that are distinct by their landscape- climatic characteristics and position within the species range. The Novosibirsk population faces a higher level of fragmentation and harsher environmental conditions than the Saratov population and is also separated from most of the species’ range by the long-standing natural barrier of the Ob river. Each of these factors can enhance the loss of genetic variation (Soulé 1973; Brussard 1984; Lawton 1993; Eckert et al. 2008) making a bottleneck or founder event more likely to occur in the Novosibirsk population compared to the Saratov population. In accordance with this expectation, both mitochondrial and microsatellite diversity were significantly lower in Novosibirsk than in Saratov. The failure to detect recent bottlenecks in either population using microsatellite loci should be interpreted with caution. First, immigration or rapid population growth can obscure the signals of past bottlenecks (Hundertmark and VanDaele 2010; Peery et al. 2012). Second, the methods used rely on the assumption of mutation-drift equilibrium (Cornuet and Luikart 1996; Garza and Williamson 2001), which was not met in our case. Deviations from Hardy–Weinberg equilibrium at many loci and the reduction in Ho compared to He in both the Novosibirsk and Saratov populations can be partly explained by fine-scale genetic substructure (Wahlund effect). Meanwhile, accelerated drift can be responsible for the significantly higher Fis-values observed in Novosibirsk compared to Saratov.

### 4.2. Genetic structure and diversity in the subterranean *E. talpinus* vs surface-dwelling voles

Gene flow between local populations/demes of subterranean rodents is believed to be limited due to their relatively restricted dispersal capabilities, compared to surface-dwelling species, as well as their dependence on specific soil types and food resources (Nevo 1999; Busch et al. 2000; Begall et al. 2007; but see Finn et al. 2022). The results of the microsatellite and D-loop-based AMOVAs, along with the DAPC, taken together, support strong genetic differentiation among *Ellobius* populations at both intermediate (dozens of kilometers, NB vs NI) and fine (several kilometers, SG vs SR and SF) spatial scales. This suggests that even small local barriers, such as water bodies and degraded steppe habitats, can cause strong genetic differentiation of this subterranean vole. In contrast, the population genetic structure of populations of the surface-dwelling vole species may be less affected by landscape heterogeneity (e.g., Ehrich et al. 2001; Gauffre et al. 2008; Adams and Hadly 2010; Stojak et al. 2016; Dominguez et al. 2021).

In small, fragmented populations, genetic variation can become quickly depleted. However, in the non-marginal Saratov population of *E. talpinus*, both microsatellite-based heterozygosity (0.76) and mitochondrial D-loop haplotype diversity (0.83) are higher than the respective average values reported for terrestrial mammals (*He* = 0.67; *Hd* = 0.58 - Saitoh 2020) and mammals in general (*He* = 0.65 - DeKort et al. 2021). Given the strong influence of phylogeny-dependent life-history traits (body size, longevity, and fertility) on genetic diversity (Eo et al. 2011; Ellegren and Galtier 2016; DeKort et al. 2021), it is more appropriate to compare the genetic pattern of the mole vole with those of phylogenetically related species, i.e. other arvicolines. The D-loop haplotype diversity detected in the Saratov population was higher than the overall mean (*Hd* ± SD: 0.63 ± 0.28) for 107 populations of eleven arvicoline species. It was also higher than the average *Hd* in ten of these species and exceeded the *Hd* of 69% of the studied populations (Table S3). Considering codominant markers, our estimate of the expected heterozygosity in the Saratov population of *E. talpinus* is close to the overall mean (*He* ± SD: 0.79 ± 0.08) calculated from the values reported for 166 local populations of 13 surface-dwelling vole species (Table S4). Intermediate or high genetic variation has previously been shown for populations of several subterranean species (*Fukomys damarensis* - Burland et al. 2001; *Ctenomys haigi* - Lacey 2001; *Nannospalax leucodon* - Karanth et al. 2004; Lin et al. 2008; Popa et al. 2014; *Bathyergus suillus* - Visser et al. 2014; *Georhychus capensis* - Visser et al. 2018). Thus, the available data do not provide support for the unambiguous impact of the subterranean lifestyle on genetic dynamics. One possible explanation is that, at certain levels of gene flow, the admixture of genotypes from different subpopulations can result in increased genetic diversity (Harrison and Hastings 1996; Hartl and Clark 1997; Amos and Harwood 1998; Alleaume-Benharira et al. 2006; Bertl et al. 2018). The strong social structure typical of many subterranean species, including mole voles, may have the same effect, minimizing the loss of genetic variation (Chesser 1991; Dobson et al. 1998; Storz 1999; Parreira and Chikhi 2015). The pattern of spatial autocorrelation revealed in *E. talpinus*, with positive r- values presented for only the first distance classes, may indicate a combination of related individuals clustered in family groups and occasional long-distance dispersal events (see Mynhardt et al. 2021 for the similar genetic pattern in *F. damarensis*). Additionally, the long generation time and iteroparity of subterranean rodents (the reported duration of reproductive life in *E. talpinus* is up to 6 years - Evdokimov 2001) can buffer losses of genetic diversity (Amos 1996; Hamrick and Godt 1996; McCoy et al. 2010; Waples et al. 2016).

### 4.3. Genetic variation in *E. talpinus*: mitochondrial vs nuclear markers

It is noteworthy that the relative polymorphism estimates for the two types of markers were different in the two populations. In Novosibirsk, the ratio He: *Hd* (0.550: 0.138 = 4.00) corresponded to the expected value based on the fact that the effective gene number for nuclear DNA is four times greater than that of the mitochondrial DNA (mtDNA), given even breeding sex ratio and equal mutation rate for both markers (Birky et al. 1983). In contrast, in the Saratov population, mitochondrial diversity was not lower but slightly higher than microsatellite diversity. This pattern is typical for terrestrial mammals (Saitoh 2021). Birky et al. (1983) attributed it to the higher mutation rate of mtDNA, but this hypothesis cannot explain the wide inter-population variations in the *He*: *Hd* ratio observed in many species (Saitoh 2021) and revealed for *E. talpinus* in this study. Alternatively or additionally, the relationship between mtDNA-based and microsatellite-based measures of genetic variation can be affected by the breeding sex ratio. High variance in female reproductive success should promote erosion of maternally inherited genes relative to biparentally inherited ones, whereas extreme polygyny should influence in the opposite direction (Briton et al. 1994; Nunney 1993; Verkuil et al. 2014). The genetic mating system of *Ellobius* species has not been well-described, but available data on family structure suggest that the northern mole vole in Saratov is a singular breeder (Bergaliev, Smorkatcheva and Rudyk, personal observation), whereas polygyny is common in the Novosibirsk population (Novikov et al. 2007; Smorkatcheva and Kuprina, personal observation). Clearly, this pattern is the opposite to what would be expected based on the high mtDNA variation compared to microsatellite variation in Saratov. Recently, a hypothesis involving male-biased dispersal has been proposed to explain the high variance of mammalian haplotype diversity (Saitoh 2021). According to this model, female philopatry promotes the divergence and spatial segregation of mitochondrial lineages. Male immigrants transfer haplotypes from other subpopulations increasing mitochondrial diversity while homogenizing the microsatellite composition of the entire recipient population. The dispersal strategy in *E. talpinus* has yet to be determined. From the visual inspection of the haplotype spatial distribution at SR (Fig. S3), no sex-specific pattern is apparent. This may be the consequence of the limited sample size and the considerable delay in natal dispersal reported for this species (Evdokimov 2001). Nevertheless, a nested subpopulation structure, where a nuclear-DNA-based population includes several mtDNA-based subpopulations, can arise even without a dispersal bias, simply due to the smaller effective number and greater sensitivity of mitochondrial genes to drift. The observed pattern of haplotype spatial distribution (Fig. S3), along with a highly ragged mismatch distribution, significantly positive Tajima’s D for the pooled SR sample, and the respective parameters for individual mitochondrial lineages are consistent with spatial expansion followed by the admixture of two or more mitochondrial subpopulations within the same weakly structured nuclear population.

## 5. CONCLUSIONS

This study represents the first characterization of the intra-population genetic structure and diversity of *Ellobius*, a highly specialized subterranean vole. The two populations studied exhibited the expected differences in genetic polymorphism: the peripheral population had low nuclear diversity and was virtually monomorphic in mtDNA, while the non-marginal population exhibited moderate nuclear and high mitochondrial variation compared to surface-dwelling voles. The spatial genetic patterns revealed in both populations appear to reflect a combination of a strong social structure, the effect of natural and anthropogenic barriers limiting dispersal opportunities, and occasional long-distance dispersal within continuous suitable habitats.

High ecological plasticity and interpopulation variations in fragmentation levels, breeding systems, and generation times (Evdokimov 2001; Novikov et al. 2007) make the northern mole vole an excellent model among subterranean species for exploring factors influencing population genetic dynamics. Based on the estimates of polymorphism revealed in two distant populations of *E. talpinus*, the selected microsatellite loci are informative for investigating kinship structure, mating system, and dispersal strategies across the species’ range. The suitability of the D-loop sequence for these purposes is less predictable, but it can be an extremely useful tool in certain populations, such as the Saratov population. Future studies of mole vole populations across a broader range could help test the roles of sex-biased dispersal, reproductive skew, social structure, and life history parameters in maintaining genetic diversity.

## Acknowledgments

This study was supported by the Russian Science Foundation (project No. 23-24-00142). The work was carried out at the Resource centers of Saint Petersburg University, “Chromas” and “Molecular and Cell Technologies”. We are grateful to Margarita M. Dymskaya, Ilya A. Volodin, Varvara R. Nikonova, Alexandra A. Panyutina, Svetlana A. Shimanovich, Alexander N. Kuznetsov, Maria S. Gribanova, Andrey O. Fedosov, Maria Y. Vdovina, Polina S. Cherepenko and Ekaterina D. Kolosova for their participation in sample collection. We are thankful to Andrey V. Tchabovsky who helped in the organization of field work, and Petr P. Strelkov and Vladimir A. Lukhtanov who provided valuable advice on data analysis.

## Authors’ contributions

Conceptualization: A.I.R., K.V.K. and A.V.S.; sample collection: A.I.R., K.V.K., A.M.B., A.V.S., E.A.N. and E.V.V.; methodology: S.A.G., K.V.K., A.I.R. and A.V.S.; resources: S.A.G., A.E.R. and E.A.N.; investigation: A.I.R., K.V.K., S.A.G., A.M.B., A.E.R. and A.V.S.; Funding acquisition: A.V.S.; writing – original draft: A.I.R., K.V.K. and A.V.S.; writing – review and editing: A.I.R., K.V.K., A.V.S., S.A.G., A.M.B., E.V.V., A.E.R. and E.A.N.

## Availability of data and materials

Sequenced fragments of the mitochondrial D-loop are available on GenBank under accession numbers: will be available after submission to the journal.

## Ethics approval

All procedures involving animals were in compliance with the national laws of the Russian Federation. The experimental protocols used in this study were approved by the Specialized Ethics Committee for Animal Research of the St. Petersburg State University (№№ 131-03- 2 and 131-03-9).

## Ethics approval consent to participate

Not applicable.

## SUPPLEMENTARY MATERIAL

**Supplementary Figure S1.**
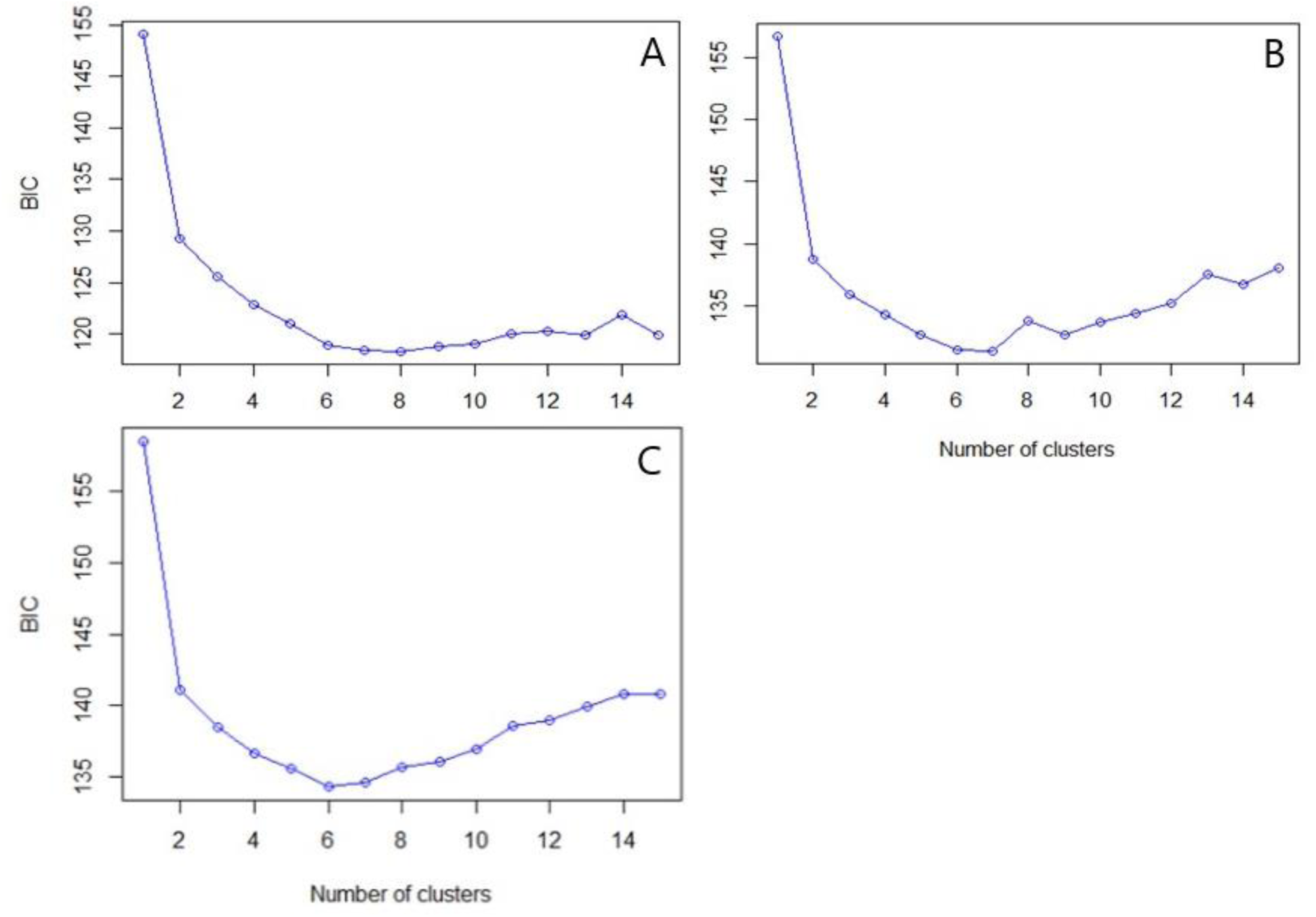
Bayesian information criteria (BIC) for different numbers of clusters with the number PCs to retain: 30 (A), 50 (B) and 70 (C) calculated for 88 individuals of *Ellobius talpinus* using nine microsatellite markers.

**Supplement Figure S2.**
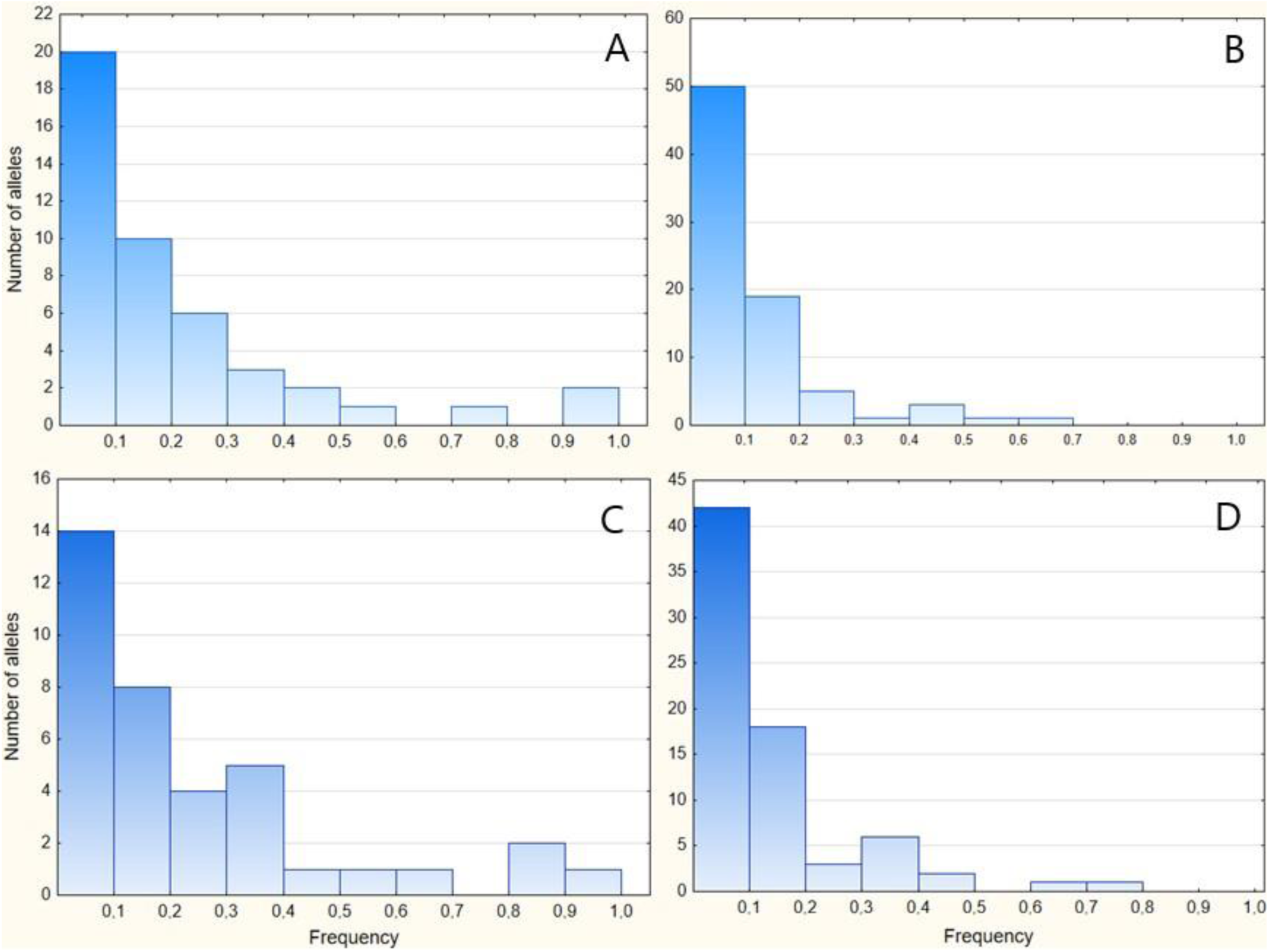
Allele frequency distributions calculated for *Ellobius talpinus* using nine microsatellite markers. A – Novosibirsk population, based on 45 alleles in 43 individuals; B – Saratov population, based on 80 alleles in 57 individuals; C – NB site, based on 37 alleles in 33 individuals, D – SR site, based on 73 alleles in 44 individuals.

**Supplementary Figure S3.**
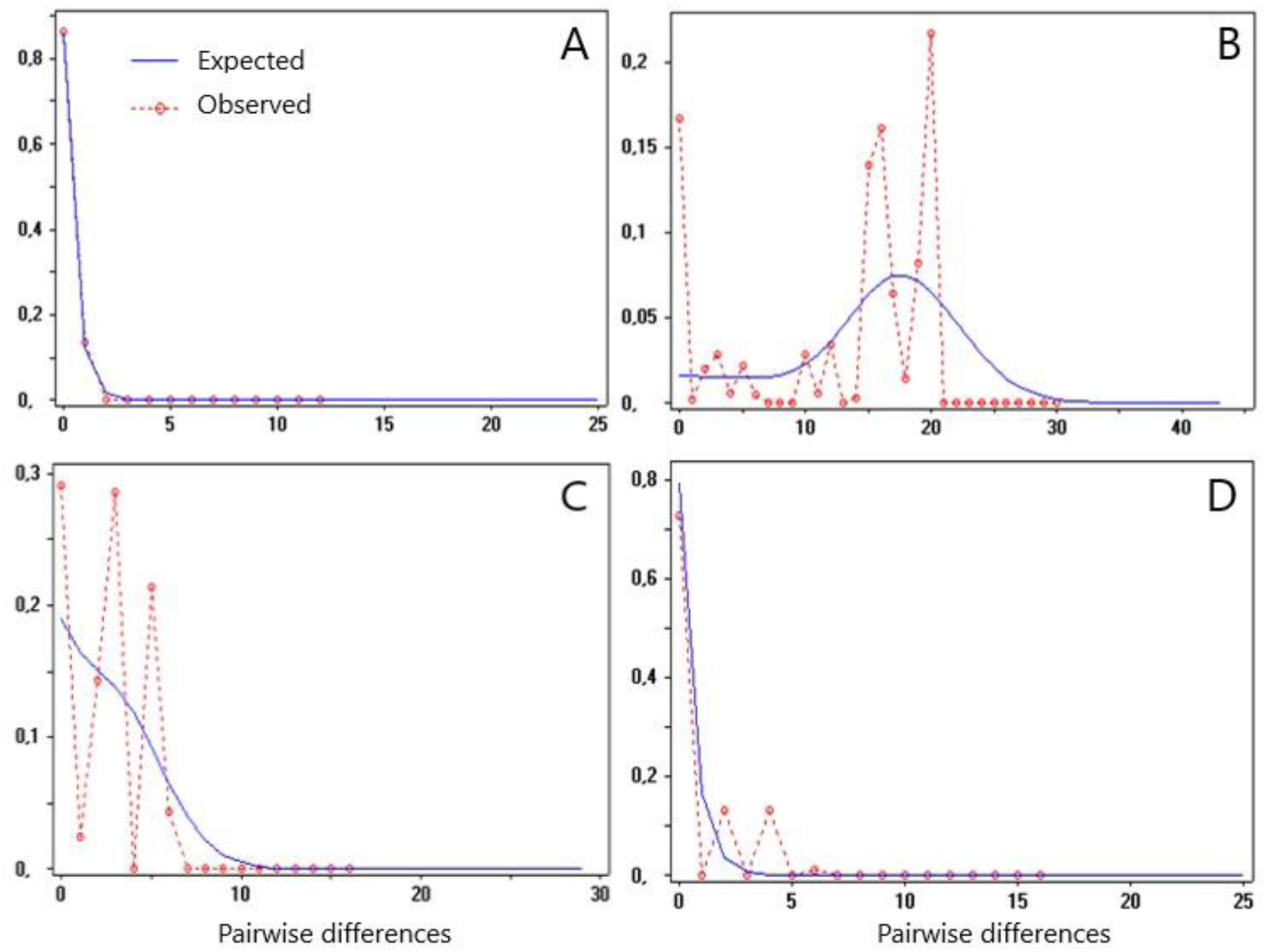
Mitochondrial mismatch distributions calculated for *Ellobius talpinus* using D-loop haplotypes. A – Novosibirsk population (n = 42); B – Saratov population (n = 65); C – Saratov population, haplogroup A (n = 21); D – Saratov population, haplogroup B (n = 14). Solid (blue) line - expected distribution under the demographic expansion model; dotted (red) line - observed distribution.

**Supplementary Figure S4.**
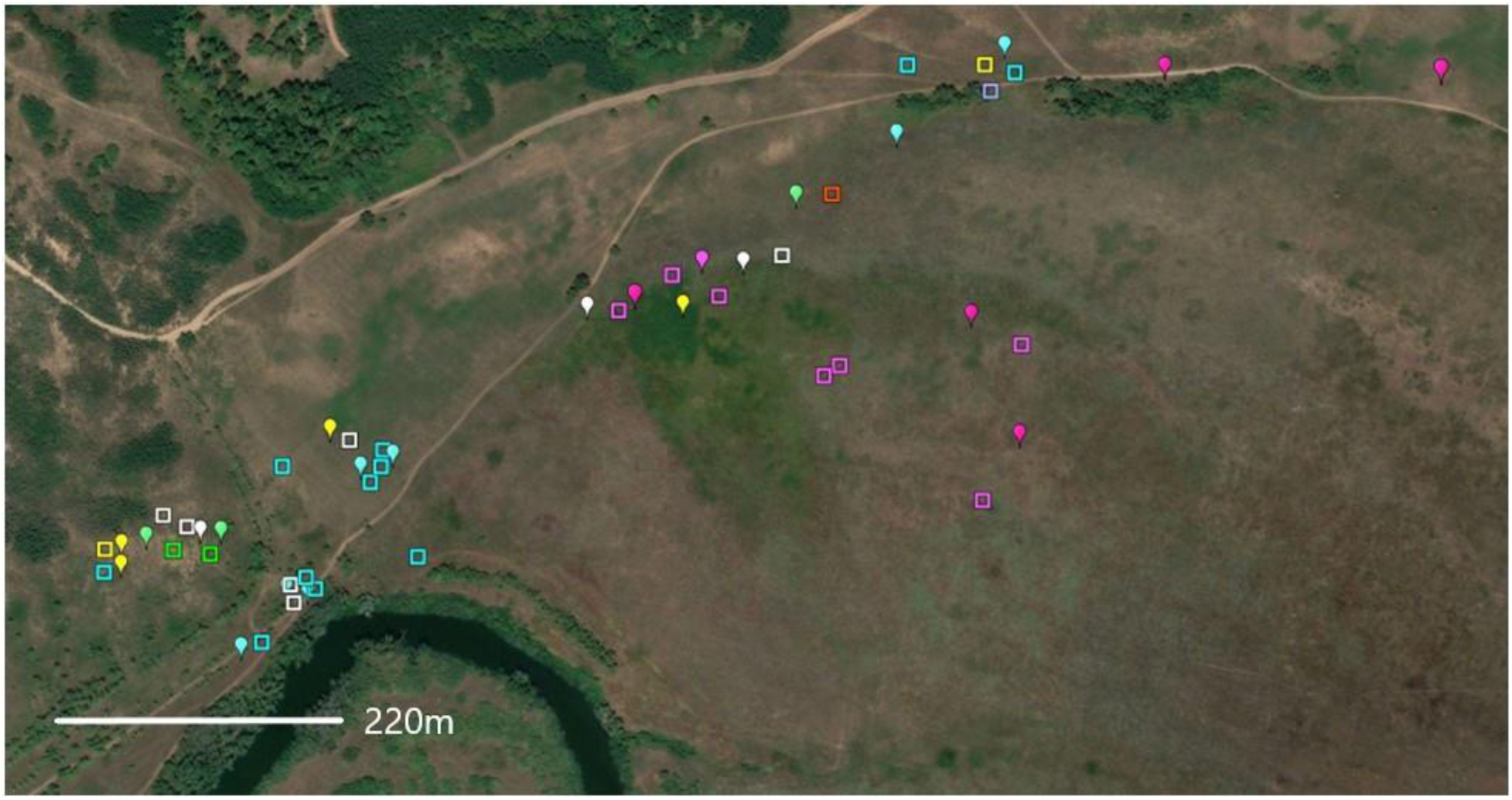
Spatial distribution of D-loop haplotypes (n = 52) of *Ellobius talpinus* within SR site. Squares represent males, drops represent females. Different colours refer to different haplotypes.

**Supplement Table S1.**
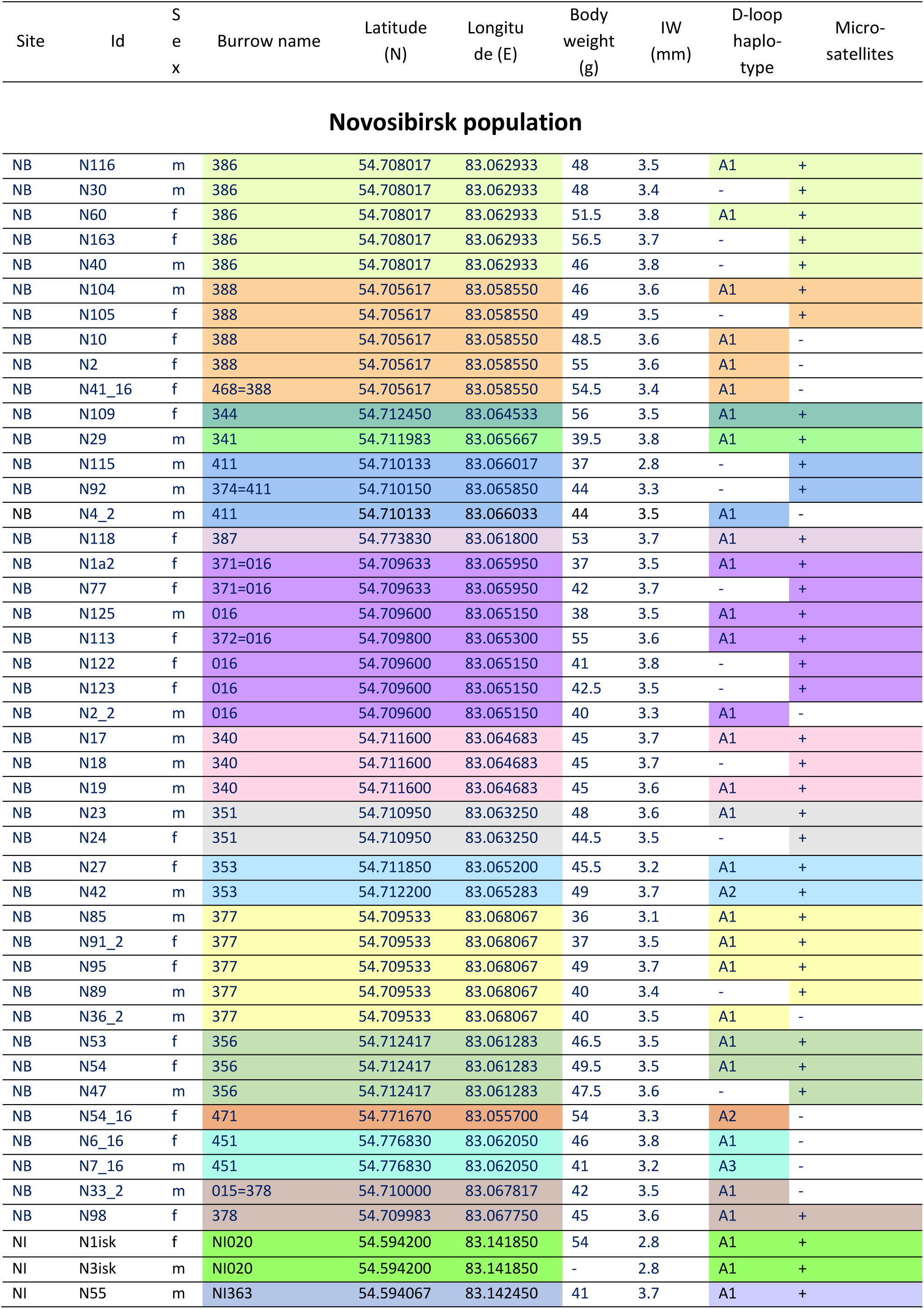
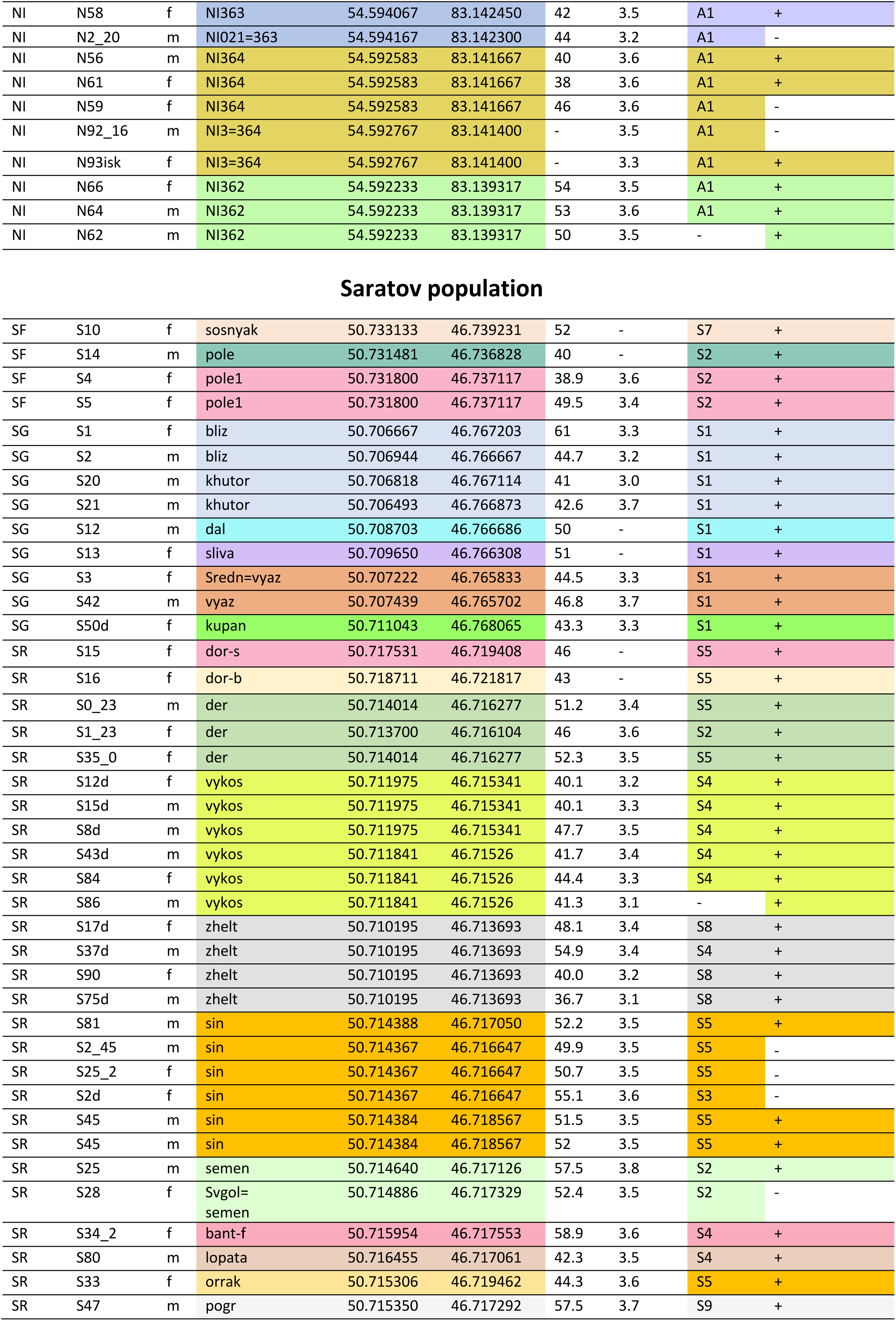
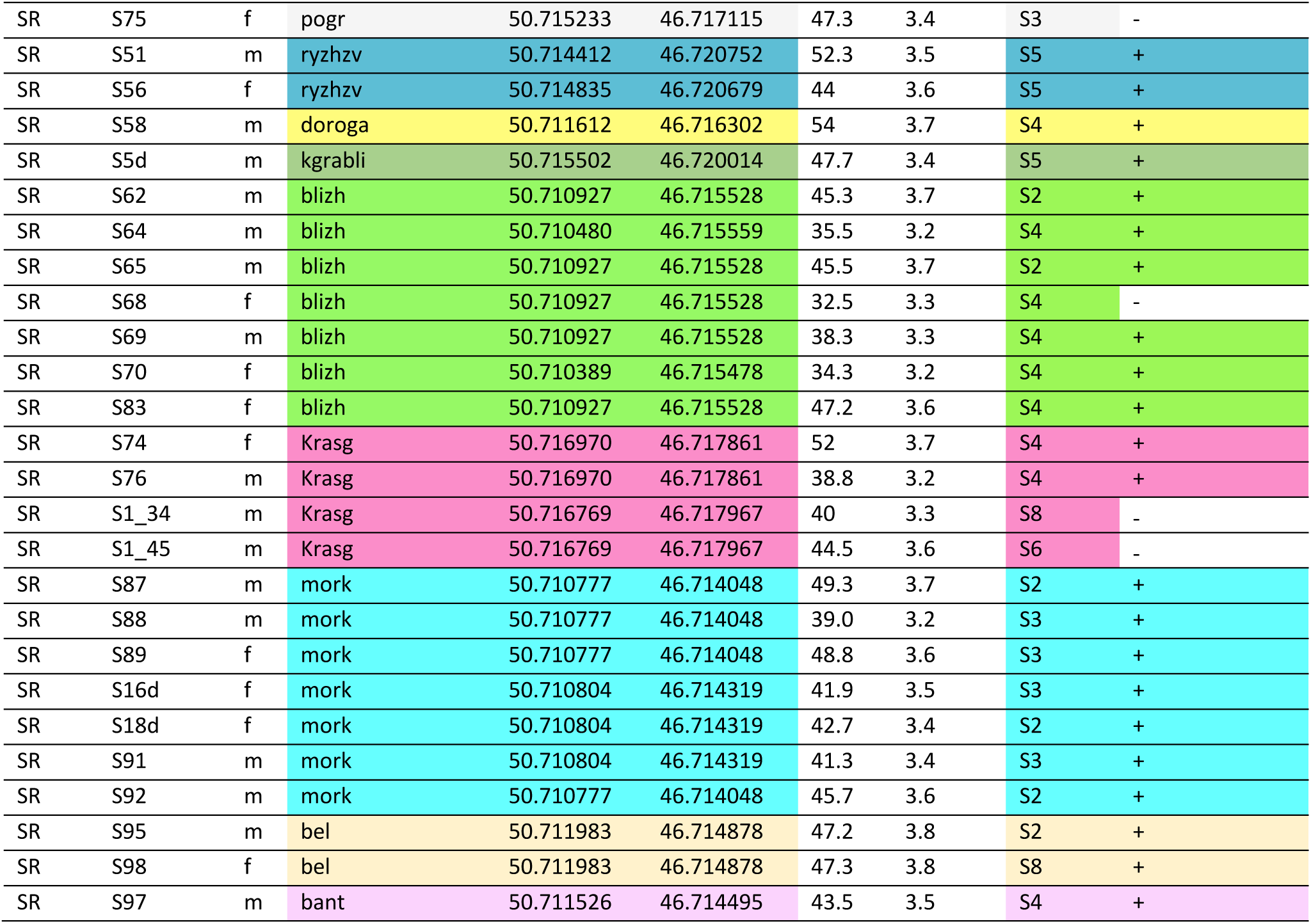
The list of samples used in the study. IW - joined width of upper incisors.

| Site | Id | Sex | Burrow name | Latitude (N) | Longitude (E) | Body weight (g) | IW (mm) | D-loop haplo-type | Micro-satellites |
| --- | --- | --- | --- | --- | --- | --- | --- | --- | --- |
| **Novosibirsk population** |  |  |  |  |  |  |  |  |  |
| NB | N116 | m | 386 | 54.708017 | 83.062933 | 48 | 3.5 | A1 | + |
| NB | N30 | m | 386 | 54.708017 | 83.062933 | 48 | 3.4 | - | + |
| NB | N60 | f | 386 | 54.708017 | 83.062933 | 51.5 | 3.8 | A1 | + |
| NB | N163 | f | 386 | 54.708017 | 83.062933 | 56.5 | 3.7 | - | + |
| NB | N40 | m | 386 | 54.708017 | 83.062933 | 46 | 3.8 | - | + |
| NB | N104 | m | 388 | 54.705617 | 83.058550 | 46 | 3.6 | A1 | + |
| NB | N105 | f | 388 | 54.705617 | 83.058550 | 49 | 3.5 | - | + |
| NB | N10 | f | 388 | 54.705617 | 83.058550 | 48.5 | 3.6 | A1 | - |
| NB | N2 | f | 388 | 54.705617 | 83.058550 | 55 | 3.6 | A1 | - |
| NB | N41_16 | f | 468=388 | 54.705617 | 83.058550 | 54.5 | 3.4 | A1 | - |
| NB | N109 | f | 344 | 54.712450 | 83.064533 | 56 | 3.5 | A1 | + |
| NB | N29 | m | 341 | 54.711983 | 83.065667 | 39.5 | 3.8 | A1 | + |
| NB | N115 | m | 411 | 54.710133 | 83.066017 | 37 | 2.8 | - | + |
| NB | N92 | m | 374=411 | 54.710150 | 83.065850 | 44 | 3.3 | - | + |
| NB | N4_2 | m | 411 | 54.710133 | 83.066033 | 44 | 3.5 | A1 | - |
| NB | N118 | f | 387 | 54.773830 | 83.061800 | 53 | 3.7 | A1 | + |
| NB | N1a2 | f | 371=016 | 54.709633 | 83.065950 | 37 | 3.5 | A1 | + |
| NB | N77 | f | 371=016 | 54.709633 | 83.065950 | 42 | 3.7 | - | + |
| NB | N125 | m | 016 | 54.709600 | 83.065150 | 38 | 3.5 | A1 | + |
| NB | N113 | f | 372=016 | 54.709800 | 83.065300 | 55 | 3.6 | A1 | + |
| NB | N122 | f | 016 | 54.709600 | 83.065150 | 41 | 3.8 | - | + |
| NB | N123 | f | 016 | 54.709600 | 83.065150 | 42.5 | 3.5 | - | + |
| NB | N2_2 | m | 016 | 54.709600 | 83.065150 | 40 | 3.3 | A1 | - |
| NB | N17 | m | 340 | 54.711600 | 83.064683 | 45 | 3.7 | A1 | + |
| NB | N18 | m | 340 | 54.711600 | 83.064683 | 45 | 3.7 | - | + |
| NB | N19 | m | 340 | 54.711600 | 83.064683 | 45 | 3.6 | A1 | + |
| NB | N23 | m | 351 | 54.710950 | 83.063250 | 48 | 3.6 | A1 | + |
| NB | N24 | f | 351 | 54.710950 | 83.063250 | 44.5 | 3.5 | - | + |
| NB | N27 | f | 353 | 54.711850 | 83.065200 | 45.5 | 3.2 | A1 | + |
| NB | N42 | m | 353 | 54.712200 | 83.065283 | 49 | 3.7 | A2 | + |
| NB | N85 | m | 377 | 54.709533 | 83.068067 | 36 | 3.1 | A1 | + |
| NB | N91_2 | f | 377 | 54.709533 | 83.068067 | 37 | 3.5 | A1 | + |
| NB | N95 | f | 377 | 54.709533 | 83.068067 | 49 | 3.7 | A1 | + |
| NB | N89 | m | 377 | 54.709533 | 83.068067 | 40 | 3.4 | - | + |
| NB | N36_2 | m | 377 | 54.709533 | 83.068067 | 40 | 3.5 | A1 | - |
| NB | N53 | f | 356 | 54.712417 | 83.061283 | 46.5 | 3.5 | A1 | + |
| NB | N54 | f | 356 | 54.712417 | 83.061283 | 49.5 | 3.5 | A1 | + |
| NB | N47 | m | 356 | 54.712417 | 83.061283 | 47.5 | 3.6 | - | + |
| NB | N54_16 | f | 471 | 54.771670 | 83.055700 | 54 | 3.3 | A2 | - |
| NB | N6_16 | f | 451 | 54.776830 | 83.062050 | 46 | 3.8 | A1 | - |
| NB | N7_16 | m | 451 | 54.776830 | 83.062050 | 41 | 3.2 | A3 | - |
| NB | N33_2 | m | 015=378 | 54.710000 | 83.067817 | 42 | 3.5 | A1 | - |
| NB | N98 | f | 378 | 54.709983 | 83.067750 | 45 | 3.6 | A1 | + |
| NI | N1isk | f | NI020 | 54.594200 | 83.141850 | 54 | 2.8 | A1 | + |
| NI | N3isk | m | NI020 | 54.594200 | 83.141850 | - | 2.8 | A1 | + |
| NI | N55 | m | NI363 | 54.594067 | 83.142450 | 41 | 3.7 | A1 | + |
| NI | N58 | f | NI363 | 54.594067 | 83.142450 | 42 | 3.5 | A1 | + |
| NI | N2_20 | m | NI021=363 | 54.594167 | 83.142300 | 44 | 3.2 | A1 | - |
| NI | N56 | m | NI364 | 54.592583 | 83.141667 | 40 | 3.6 | A1 | + |
| NI | N61 | f | NI364 | 54.592583 | 83.141667 | 38 | 3.6 | A1 | + |
| NI | N59 | f | NI364 | 54.592583 | 83.141667 | 46 | 3.6 | A1 | - |
| NI | N92_16 | m | NI3=364 | 54.592767 | 83.141400 | - | 3.5 | A1 | - |
| NI | N93isk | f | NI3=364 | 54.592767 | 83.141400 | - | 3.3 | A1 | + |
| NI | N66 | f | NI362 | 54.592233 | 83.139317 | 54 | 3.5 | A1 | + |
| NI | N64 | m | NI362 | 54.592233 | 83.139317 | 53 | 3.6 | A1 | + |
| NI | N62 | m | NI362 | 54.592233 | 83.139317 | 50 | 3.5 | - | + |
| Saratov population |  |  |  |  |  |  |  |  |  |
| SF | S10 | f | sosnyak | 50.733133 | 46.739231 | 52 | - | S7 | + |
| SF | S14 | m | pole | 50.731481 | 46.736828 | 40 | - | S2 | + |
| SF | S4 | f | pole1 | 50.731800 | 46.737117 | 38.9 | 3.6 | S2 | + |
| SF | S5 | f | pole1 | 50.731800 | 46.737117 | 49.5 | 3.4 | S2 | + |
| SG | S1 | f | bliz | 50.706667 | 46.767203 | 61 | 3.3 | S1 | + |
| SG | S2 | m | bliz | 50.706944 | 46.766667 | 44.7 | 3.2 | S1 | + |
| SG | S20 | m | khutor | 50.706818 | 46.767114 | 41 | 3.0 | S1 | + |
| SG | S21 | m | khutor | 50.706493 | 46.766873 | 42.6 | 3.7 | S1 | + |
| SG | S12 | m | dal | 50.708703 | 46.766686 | 50 | - | S1 | + |
| SG | S13 | f | sliva | 50.709650 | 46.766308 | 51 | - | S1 | + |
| SG | S3 | f | Sredn=vyaz | 50.707222 | 46.765833 | 44.5 | 3.3 | S1 | + |
| SG | S42 | m | vyaz | 50.707439 | 46.765702 | 46.8 | 3.7 | S1 | + |
| SG | S50d | f | kupan | 50.711043 | 46.768065 | 43.3 | 3.3 | S1 | + |
| SR | S15 | f | dor-s | 50.717531 | 46.719408 | 46 | - | S5 | + |
| SR | S16 | f | dor-b | 50.718711 | 46.721817 | 43 | - | S5 | + |
| SR | S0_23 | m | der | 50.714014 | 46.716277 | 51.2 | 3.4 | S5 | + |
| SR | S1_23 | f | der | 50.713700 | 46.716104 | 46 | 3.6 | S2 | + |
| SR | S35_0 | f | der | 50.714014 | 46.716277 | 52.3 | 3.5 | S5 | + |
| SR | S12d | f | vykos | 50.711975 | 46.715341 | 40.1 | 3.2 | S4 | + |
| SR | S15d | m | vykos | 50.711975 | 46.715341 | 40.1 | 3.3 | S4 | + |
| SR | S8d | m | vykos | 50.711975 | 46.715341 | 47.7 | 3.5 | S4 | + |
| SR | S43d | m | vykos | 50.711841 | 46.71526 | 41.7 | 3.4 | S4 | + |
| SR | S84 | f | vykos | 50.711841 | 46.71526 | 44.4 | 3.3 | S4 | + |
| SR | S86 | m | vykos | 50.711841 | 46.71526 | 41.3 | 3.1 | - | + |
| SR | S17d | f | zhelt | 50.710195 | 46.713693 | 48.1 | 3.4 | S8 | + |
| SR | S37d | m | zhelt | 50.710195 | 46.713693 | 54.9 | 3.4 | S4 | + |
| SR | S90 | f | zhelt | 50.710195 | 46.713693 | 40.0 | 3.2 | S8 | + |
| SR | S75d | m | zhelt | 50.710195 | 46.713693 | 36.7 | 3.1 | S8 | + |
| SR | S81 | m | sin | 50.714388 | 46.717050 | 52.2 | 3.5 | S5 | + |
| SR | S2_45 | m | sin | 50.714367 | 46.716647 | 49.9 | 3.5 | S5 | - |
| SR | S25_2 | f | sin | 50.714367 | 46.716647 | 50.7 | 3.5 | S5 | - |
| SR | S2d | f | sin | 50.714367 | 46.716647 | 55.1 | 3.6 | S3 | - |
| SR | S45 | m | sin | 50.714384 | 46.718567 | 51.5 | 3.5 | S5 | + |
| SR | S45 | m | sin | 50.714384 | 46.718567 | 52 | 3.5 | S5 | + |
| SR | S25 | m | semen | 50.714640 | 46.717126 | 57.5 | 3.8 | S2 | + |
| SR | S28 | f | Svgol=semen | 50.714886 | 46.717329 | 52.4 | 3.5 | S2 | - |
| SR | S34_2 | f | bant-f | 50.715954 | 46.717553 | 58.9 | 3.6 | S4 | + |
| SR | S80 | m | lopata | 50.716455 | 46.717061 | 42.3 | 3.5 | S4 | + |
| SR | S33 | f | orrak | 50.715306 | 46.719462 | 44.3 | 3.6 | S5 | + |
| SR | S47 | m | pogr | 50.715350 | 46.717292 | 57.5 | 3.7 | S9 | + |
| SR | S75 | f | pogr | 50.715233 | 46.717115 | 47.3 | 3.4 | S3 | - |
| SR | S51 | m | ryzhzv | 50.714412 | 46.720752 | 52.3 | 3.5 | S5 | + |
| SR | S56 | f | ryzhzv | 50.714835 | 46.720679 | 44 | 3.6 | S5 | + |
| SR | S58 | m | doroga | 50.711612 | 46.716302 | 54 | 3.7 | S4 | + |
| SR | S5d | m | kgrabli | 50.715502 | 46.720014 | 47.7 | 3.4 | S5 | + |
| SR | S62 | m | blizh | 50.710927 | 46.715528 | 45.3 | 3.7 | S2 | + |
| SR | S64 | m | blizh | 50.710480 | 46.715559 | 35.5 | 3.2 | S4 | + |
| SR | S65 | m | blizh | 50.710927 | 46.715528 | 45.5 | 3.7 | S2 | + |
| SR | S68 | f | blizh | 50.710927 | 46.715528 | 32.5 | 3.3 | S4 | - |
| SR | S69 | m | blizh | 50.710927 | 46.715528 | 38.3 | 3.3 | S4 | + |
| SR | S70 | f | blizh | 50.710389 | 46.715478 | 34.3 | 3.2 | S4 | + |
| SR | S83 | f | blizh | 50.710927 | 46.715528 | 47.2 | 3.6 | S4 | + |
| SR | S74 | f | Krasg | 50.716970 | 46.717861 | 52 | 3.7 | S4 | + |
| SR | S76 | m | Krasg | 50.716970 | 46.717861 | 38.8 | 3.2 | S4 | + |
| SR | S1_34 | m | Krasg | 50.716769 | 46.717967 | 40 | 3.3 | S8 | - |
| SR | S1_45 | m | Krasg | 50.716769 | 46.717967 | 44.5 | 3.6 | S6 | - |
| SR | S87 | m | mork | 50.710777 | 46.714048 | 49.3 | 3.7 | S2 | + |
| SR | S88 | m | mork | 50.710777 | 46.714048 | 39.0 | 3.2 | S3 | + |
| SR | S89 | f | mork | 50.710777 | 46.714048 | 48.8 | 3.6 | S3 | + |
| SR | S16d | f | mork | 50.710804 | 46.714319 | 41.9 | 3.5 | S3 | + |
| SR | S18d | f | mork | 50.710804 | 46.714319 | 42.7 | 3.4 | S2 | + |
| SR | S91 | m | mork | 50.710804 | 46.714319 | 41.3 | 3.4 | S3 | + |
| SR | S92 | m | mork | 50.710777 | 46.714048 | 45.7 | 3.6 | S2 | + |
| SR | S95 | m | bel | 50.711983 | 46.714878 | 47.2 | 3.8 | S2 | + |
| SR | S98 | f | bel | 50.711983 | 46.714878 | 47.3 | 3.8 | S8 | + |
| SR | S97 | m | bant | 50.711526 | 46.714495 | 43.5 | 3.5 | S4 | + |

**Supplementary Table S2.** Summary of microsatellite variation in *Ellobius talpinus* from two populations and two sites with sample sizes >20. A - number of alleles per locus, Ar - rarefied allelic richness (based on the smallest population sample size for NB); Ho (±SE) - observed heterozygosity, uHe (±SE) - unbiased expected heterozygosity; HW - results of Hardy-Weinberg tests. The estimated frequencies of null alleles at each locus are shown. An asterisk indicates significant departure from HWE after Holm’s sequential Bonferroni correction.

|  | Population | n | A | Ar | Ho | uHe | Fis | H-W<br>p-value | Null allele<br>frequency |
| --- | --- | --- | --- | --- | --- | --- | --- | --- | --- |
| Mar076 | NB | 33 | 5 | 4.99 | 0.424 | 0.712 | 0.408 | <0.001* | 0.197 |
|  | Novosibirsk, total | 43 | 6 | 4.50 | 0.442 | 0.774 | 0.432 | <0.001* | 0.207 |
|  | SR | 44 | 6 | 5.93 | 0.773 | 0.749 | - | 0.346 | -0.027 |
|  | Saratov, total | 57 | 7 | 4.64 | 0.737 | 0.778 | 0.033 | 0.043 | 0.024 |
|  |  |  |  |  |  |  | 0.062 |  |  |
| Mar003 | NB | 33 | 2 | 2.00 | 0.212 | 0.282 | 0.251 | 0.190 | 0.099 |
|  | Novosibirsk, total | 42 | 4 | 3.00 | 0.214 | 0.457 | 0.534 | <0.001* | 0.222 |
|  | SR | 44 | 7 | 6.94 | 0.705 | 0.770 | 0.086 | 0.005* | 0.025 |
|  | Saratov, total | 57 | 8 | 4.31 | 0.649 | 0.805 | 0.195 | <0.001* | 0.086 |
| Mar063 | NB | 33 | 3 | 3.00 | 0.333 | 0.589 | 0.438 | <0.001* | 0.189 |
|  | Novosibirsk, total | 43 | 4 | 4.00 | 0.372 | 0.609 | 0.392 | <0.001* | 0.174 |
|  | SR | 44 | 8 | 7.96 | 0.659 | 0.816 | 0.194 | 0.023 | 0.089 |
|  | Saratov, total | 57 | 9 | 4.99 | 0.632 | 0.841 | 0.195 | 0.001* | 0.120 |
| Cg16E5 | NB | 26 | 3 | 3.00 | 0.231 | 0.212 | 0.091 | 1.000 | -0.121 |
|  | Novosibirsk, total | 36 | 3 | 3.00 | 0.167 | 0.157 | - | 1.000 | -0.086 |
|  | SR | 44 | 8 | 6.95 | 0.591 | 0.594 | 0.063 | 0.452 | -0.033 |
|  | Saratov, total | 57 | 8 | 5.32 | 0.614 | 0.675 | 0.005 | 0.090 | 0.028 |
|  |  |  |  |  |  |  | 0.092 |  |  |
| Cg16H64 | NB | 33 | 9 | 8.53 | 0.576 | 0.781 | 0.266 | 0.001* | 0.125 |
|  | Novosibirsk, total | 41 | 14 | 8.26 | 0.634 | 0.832 | 0.240 | 0.001* | 0.119 |
|  | SR | 42 | 17 | 15.1 | 0.881 | 0.897 | 0.018 | 0.215 | 0.003 |
|  | Saratov, total | 55 | 20 | 8.37 | 0.855 | 0.918 | 0.069 | 0.027 | 0.030 |
| Cg5G6 | NB | 31 | 1 | 1.00 | 0 | 0 | na | na | na |
|  | Novosibirsk, total | 41 | 1 | 1.50 | 0 | 0 | na | na | na |
|  | SR | 44 | 6 | 5.01 | 0.386 | 0.445 | 0.134 | 0.195 | 0.051 |
|  | Saratov, total | 57 | 6 | 3.00 | 0.316 | 0.569 | 0.447 | <0.001* | 0.194 |
| Mb24n | NB | 33 | 3 | 3.00 | 0.273 | 0.485 | 0.441 | 0.005* | 0.193 |
|  | Novosibirsk, total | 42 | 3 | 3.00 | 0.262 | 0.615 | 0.577 | <0.001* | 0.262 |
|  | SR | 44 | 3 | 3.84 | 0.568 | 0.614 | 0.076 | 0.532 | 0.039 |
|  | Saratov, total | 57 | 4 | 2.95 | 0.579 | 0.619 | 0.066 | 0.577 | 0.034 |
| Mb03n | NB | 32 | 7 | 6.78 | 0.500 | 0.786 | 0.367 | 0.001* | 0.176 |
|  | Novosibirsk, total | 42 | 7 | 6.39 | 0.452 | 0.789 | 0.429 | <0.001 | 0.206 |
|  | SR | 44 | 8 | 7.74 | 0.614 | 0.756 | 0.190 | 0.014 | 0.091 |
|  | Saratov, total | 57 | 8 | 5.25 | 0.596 | 0.769 | 0.225 | 0.004* | 0.109 |
| MSMM2 | NB | 33 | 4 | 4.00 | 0.485 | 0.706 | 0.317 | 0.008 | 0.142 |
|  | Novosibirsk, total | 43 | 4 | 4.00 | 0.322 | 0.537 | 0.292 | 0.007 | 0.122 |
|  | SR | 42 | 9 | 8.75 | 0.786 | 0.839 | 0.064 | 0.104 | 0.028 |
|  | Saratov, total | 55 | 10 | 4.58 | 0.581 | 0.688 | 0.075 | 0.012 | 0.034 |

**Supplementary Table S3.**
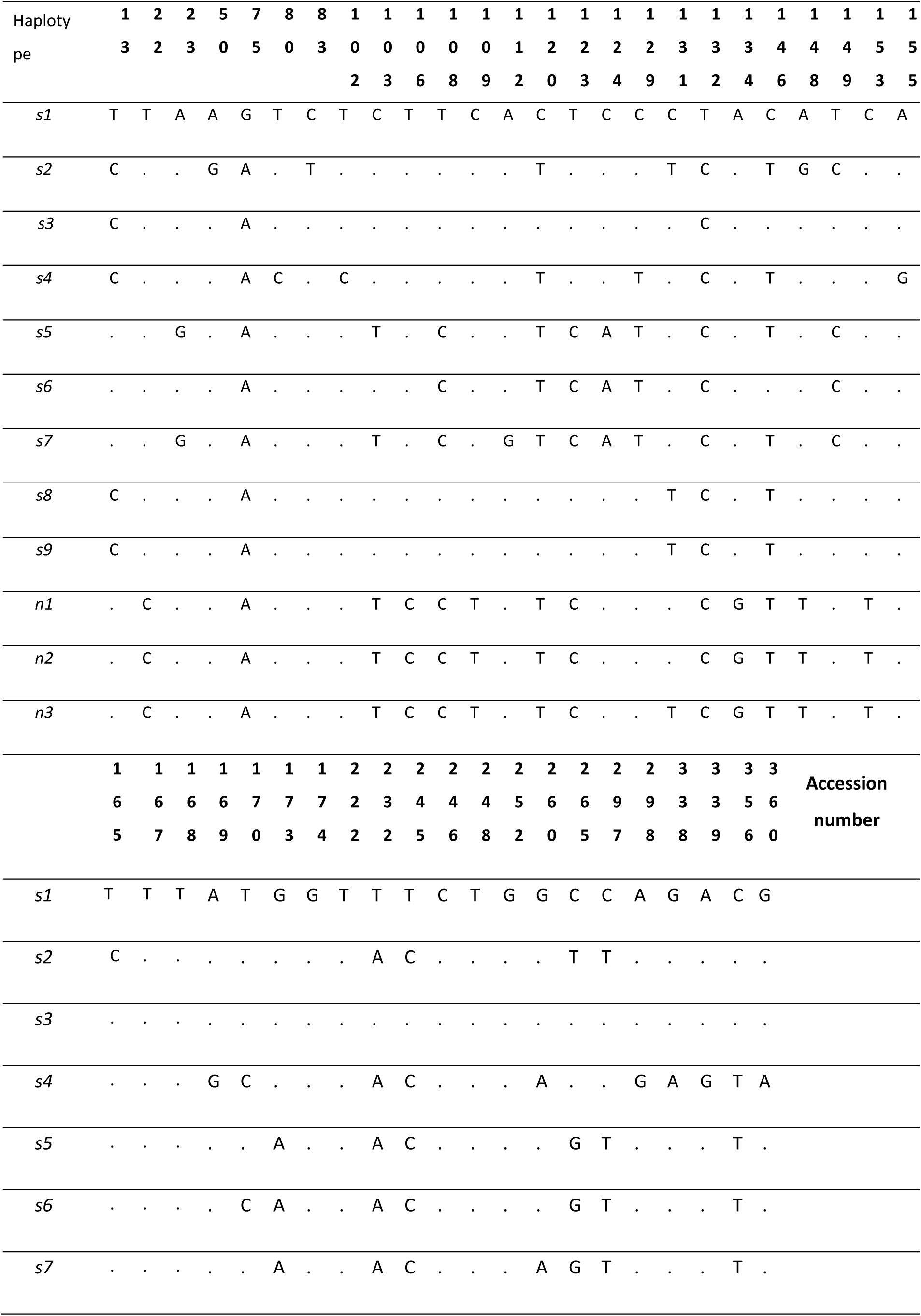
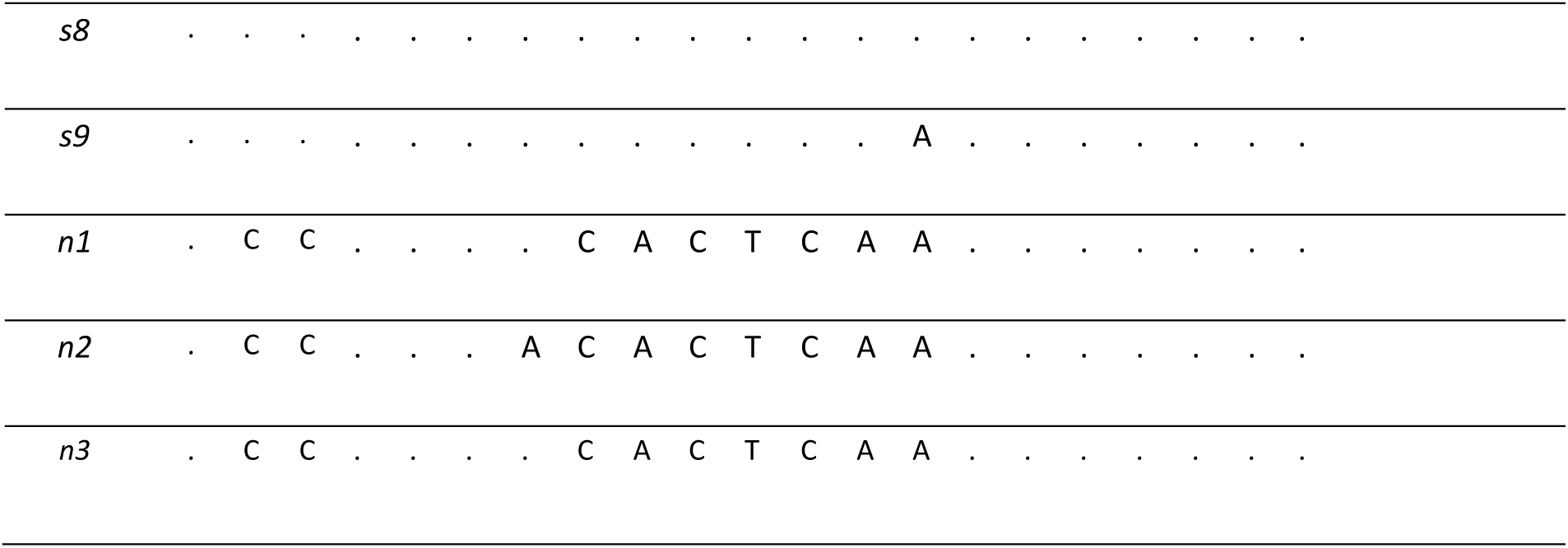
Positions of the variable nucleotides that define the haplotypes detected in the Saratov (*s1-s9*) and Novosibirsk (*n1-n3*) populations of *E. talpinus*. Haplotype *s1* is shown as the reference; dots indicate nucleotides identical to the reference.

